# Uncovering dynamic human brain phase coherence networks

**DOI:** 10.1101/2024.11.15.623830

**Authors:** Anders S. Olsen, Anders Brammer, Patrick M. Fisher, Morten Mørup

## Abstract

Complex cognitive functions rely on coordinated communication between distributed brain regions, yet capturing these interactions as they evolve over time remains challenging. Traditional analyses of functional brain connectivity largely rely on correlations in signal amplitude, which are sensitive to noise and artifacts such as head motion. Here, we introduce a mixture modeling approach that focuses on the phase of brain signals, allowing dynamic patterns of large-scale synchronization in brain phase coherence networks to be studied directly and in their entirety. We lay the mathematical and conceptual groundwork for phase modeling and introduce the complex angular central Gaussian mixture model, providing a principled way to analyze phase-based interactions across the brain. Applied to fMRI data, the model identifies recurring states of brain-wide synchronized activity that reliably distinguish cognitive tasks and generalize across previously unseen individuals, without requiring any task labels during training. These results show that modeling signal phase offers a clean and informative view of brain synchronization dynamics, opening new avenues for studying large-scale neural coordination.

**Significance statement:** Understanding how the human brain coordinates activity across distant regions is central to explaining cognition and behavior. Most existing approaches study these interactions by tracking changes in signal strength, which can be strongly affected by non-neural artifacts. Here, we focus instead on the phase relationships between brain signals: How brain regions synchronize their oscillations forming dynamic phase coherence networks. We introduce a flexible and principled mixture modeling framework to capture these patterns directly and reveal recurring states of brain-wide synchronized activity consistent across individuals. This approach uncovers meaningful differences between diverse cognitive tasks, without requiring task labels during training. By emphasizing signal phase rather than amplitude, our method offers a complementary robust and interpretable way to study dynamic brain synchronization patterns.

## Introduction

Synchronization is a fundamental concept in science, observed across diverse fields from coherent light in lasers, anti-coherence in acoustic noise suppression, and the circadian rhythm aligning with the dark-light cycle of the environment. In neuroscience, synchronization is crucial across multiple scales: Neuron-to-neuron communication often occurs through synchronized bursts of firing (Coombes and Bressloff 2005; Maeda, Robinson, and Kawana 1995) and synchronized mesoscale activity within cortical neuronal columns generates electrical signals that can be detected on the scalp (Kirschstein and Köhling 2009). At the macroscale, synchronization underpins functional connectivity across brain areas, providing a framework for understanding cognitive functions and behaviors (Bola and Sabel 2015; Uhlhaas et al. 2009; Womelsdorf et al. 2007). As such, accurate models of phase coherence, the quantification of synchronization, in multi-channel brain data are essential to capture the complexity of brain-wide connectivity patterns, key to understanding the orchestrated dynamics within and across neural networks.

Traditionally, connectivity between regional neuroimaging signals such as those from blood oxygen level dependent functional magnetic resonance imaging (BOLD fMRI) has been measured using the Pearson correlation coefficient (Biswal et al. 1995), which captures the linear relationship between signals (Bastos and Schoffelen 2016; H. E. Wang et al. 2014). Estimating time-varying correlation coefficients requires temporal windowing of input signals, potentially decreasing stationarity of low-frequency components and leading to challenges in determining, e.g., the optimal window length, function, and stride (Allen et al. 2014; Leonardi and Van De Ville 2015; S. F. Nielsen et al. 2017; Preti, Bolton, and Van De Ville 2017), and has been suggested to be an unstable approach to identifying dynamic functional connectivity (Choe et al. 2017; Hindriks et al. 2016; Laumann, Snyder, Mitra, et al. 2017; Liégeois, Laumann, et al. 2017), see (Laumann, Snyder, and Gratton 2024) for a recent review. Directly modeling the time-series using Gaussian mixture models (GMM) or hidden Markov models offer window-free alternatives (Vidaurre, Smith, and Woolrich 2017), however, intensity-based time-series methods are vulnerable to amplitude-related noise such as head motion (Chen, Rubinov, and Chang 2017). Instead, assessing phase coherence (a.k.a. phase coupling) between brain signals, focusing exclusively on oscillations and discarding (co-)amplitude information, offers a frame-wise connectivity proxy free from area-specific amplitude-related noise uninformative of connectivity (Glerean et al. 2012).

Phase estimation involves converting signals to the complex domain, typically via the Hilbert transform, which translates the temporal evolution of a signal *x*(*t*) into a revolution around the unit circle *z*(*t*) = *a*(*t*) e^*iθ*(*t*)^ (Fig. 1A). The phase *θ*(*t*) is the angle to the real axis, and co-oscillation between regions is reflected by phase differences assessed at each data sample, i.e., *θ*_*j*_(*t*) − *θ*_*j*_′ (*t*) for regions *j* and *j*^′^. Since the phase and phase difference is a circular quantity with period 2*π* and therefore difficult to model arithmetically, most previous fMRI-studies involving phase coherence have focused exclusively on cosine phase coherence cos (*θ*_*j*_(*t*) − *θ*_*j*_′ (*t*)) (Cabral et al. 2017; Glerean et al. 2012; Honari, Choe, and Lindquist 2021; ML et al. 2020; Zarghami, Hossein-Zadeh, and Bahrami 2020), discarding the imaginary dimension (the sine), which holds potentially valuable information about phase alignment. In contrast, the sine is favored in sensor-level electroencephalography data due to volume conduction (Nolte et al. 2004). Some studies have focused exclusively on the leading eigenvector of the cosine phase coherence matrix, known as Leading Eigenvector Dynamics Analysis (LEiDA) (Cabral et al. 2017), further explicitly discarding a portion of the data variance within the cosine projection of phase coherence.

**Figure 1.**
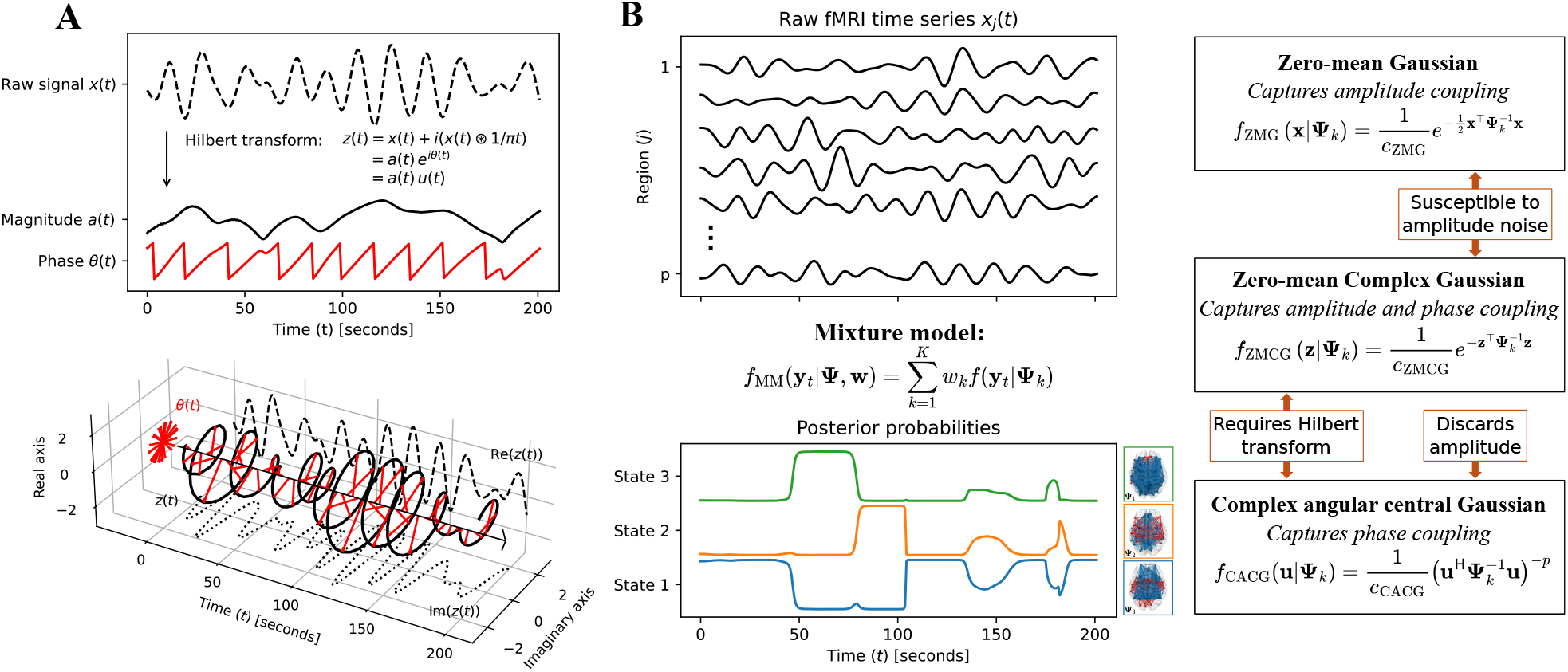
Mixture modeling of fMRI signals using either their raw values, phase or phase-amplitude representations. **A**): The Hilbert transform can be used to create the analytic signal *z*(*t*), which decomposes a time-series *x*(*t*) into phase *θ*(*t*) and amplitude *a*(*t*). The rectangular representation of phase, discarding amplitude and normalizing by dimensionality, is 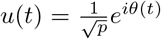. **B**) Mixture modeling of fMRI data can be approached using either their Euclidean representation (zero-mean Gaussian), or through the Hilbert transform providing a phase-amplitude representation (complex zero-mean Gaussian). Alternatively, the amplitude can be discarded completely and the phase modeled using the complex angular central Gaussian (CACG) distribution. *c*_(*·*)_ is a normalization constant (see Methods for full densities).

In this paper, we show that interregional phase coherence is very easily modeled when considering the full rectangular form 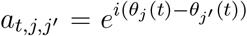. The full phase coherence map **A**_*t*_ ∈ ℂ^*p*×*p*^ for each volume *t* is rank-1, i.e., 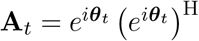, where (·)^H^ is the complex conjugate transpose, rendering modeling a pool of instantaneous complex-valued phase vectors 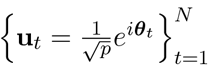 straightforward without the need for dimensionality reduction. Importantly, phase vectors are equi-norm and sign-invariant. Such data are embedded in the complex projective hyperplane ℂ ℙ*p*−1. The Gaussian analogue on this manifold is the complex angular central Gaussian (CACG) distribution (Methods) (Kent 1997; Tyler 1987), which is parameterized by a full covariance matrix. However, macroscale brain function has routinely been posited to be low rank (Shine et al. 2019; Rué-Queralt et al. 2021) and with increasing spatial resolution the number of estimated pairwise covariances increases quadratically. Therefore, a rank-*r* reparameterization, where *r* ≤ *p*, can be applied to explicitly control the number of estimated parameters, deriving a more parsimonious and interpretable representation of brain connectivity.

Here we propose a new dynamic modeling framework for the detailed characterization of phase coherence networks using probabilistic mixture modeling with CACG distributions as mixture components (Fig. 1B). We contrast our phase coherence modeling with traditional time-series modeling via the Gaussian distribution, phase-amplitude modeling via the complex Gaussian distribution, and existing K-means approaches for phase coherence, including LEiDA (see Supporting Table S1 for an overview of all models). All mixture components are parameterized by a covariance matrix, the rank of which we show is important for characterizing the complexity of dynamic states in the data. We apply our mixture models, which are trained without the use of labels, to task fMRI data from the Human Connectome project (David C. Van Essen et al. 2013), showcasing how the CACG mixture model learns meaningful phase coherence components reflective of cognitive states. When applied to resting-state fMRI, the model recovers relatively few, complex network patterns. In short, phase modeling with CACG mixtures offers a principled, robust, and interpretable unsupervised toolkit for mapping dynamic brain phase coherence networks. All models have been implemented in our openly available Python toolbox “Phase Coherence Mixture Modeling” (PCMM) (github.com/anders-s-olsen/PCMM).

## Results

We first establish the advantage of modeling phase coherence in its entirety using the complex domain, accounting for both the cosine and sine. We subsequently introduce the CACG distribution as the Gaussian analogue for multivariate phase data embedded in the complex projective hyperplane and fit a CACG mixture model to task-based and resting-state fMRI data from 255 unrelated subjects from the Human Connectome Project. We showcase the impact of component complexity on model estimates and task classification accuracy, and the benefits of phase coherence modeling compared to amplitude- and phase-amplitude modeling.

### The complex domain is required for modeling phase coherence in its entirety

To estimate phase coherence, signals are first converted to the complex domain via the Hilbert transform (Fig. 1A), wherein the amplitude is discarded and only the complex-valued instantaneous phase vectors 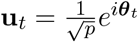 retained. A model can then be fitted to a pool of phase vectors, explaining consistent phase variations. As illustrated in Fig. 2, modeling only the real part (cosine) of the complex phase vectors as suggested in, among others, (Cabral et al. 2017; Glerean et al. 2012), results in phase shifts of *θ* and 2*π* − *θ*, i.e., reflections around the real axis, becoming indistinguishable. Similarly, modeling only the imaginary part of coherence (Nolte et al. 2004) results in signals that are in phase becoming indistinguishable from signals in antiphase (reflection around the imaginary axis). Only the complex representation can account for all consistent phase deviations as opposed to random, uncorrelated signals. Consequently, the full complex representation is needed to account for all possible types of phase coherence and previous methods focusing on, e.g., only cosine or sine coherence are inherently limited in their representational value.

**Figure 2.**
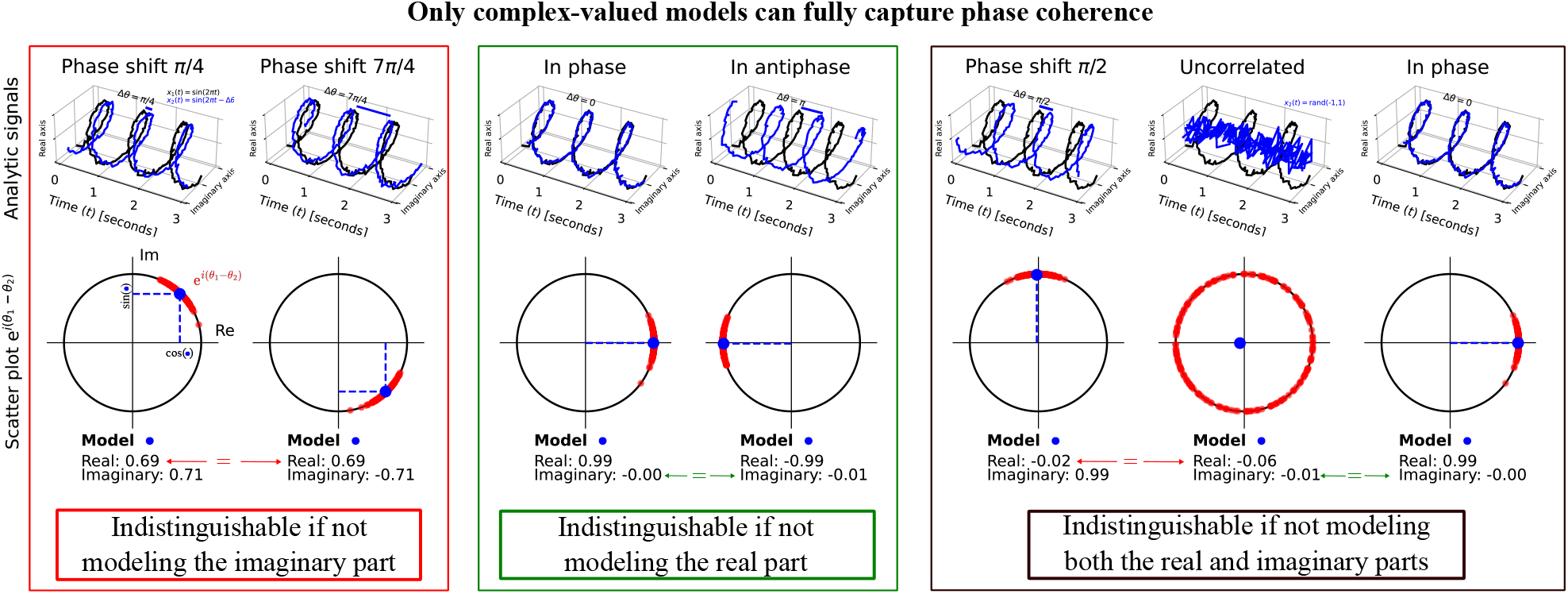
Modeling phase coherence in its entirety. Two oscillatory signals can be compared by examining their phases, which indicate the position of each sample within the oscillation cycle. The phase difference of such signals indicates their average displacement (phase shift). Gathering many phase time series, e.g., from fMRI data, enables modeling the coherence between all region pairs. However, previous models have resorted to only modeling the cosine of pairwise phase differences, meaning that phase shifts *θ* and 2*π* − *θ* (red box) would lead to the same model estimate. Likewise, phase shifts of ± *π/*2 cannot be distinguished from uncorrelated signals (black box) using cosine models. Similarly, modeling only the sine, corresponding to the imaginary part, fails to distinguish in-phase from antiphasic signals (green box) and in-phase from uncorrelated signals (black box). These issues are resolved when modeling both cosine and sine in a combined complex-valued representation.

To confirm the practical implications of not modeling the full complex-valued phase coherence, we generated synthetic data from *K* = 2 true components with consistent phase shifts and varying noise levels in *p* = 3 dimensions (Supporting Fig. S1). As expected, mixture models for the complex-valued phase coherence, including the CACG model, best recovered the original data distribution.

### Rank-1 component covariances are insufficient for modeling anisotropic data

Multivariate brain data is anisotropic over time; i.e., two adjacent brain areas are likely to have similar neuroimaging time-series while a third brain area may have a time-series unrelated to the first two (Fig. 3A). In this case, the data distribution tends to a low-rank manifold, e.g., a hyperplane in the Euclidean vector space or a great circle for phase data on the projective hyperplane. Importantly, the rank of the subspace can fall anywhere between one and the data’s dimensionality.

**Figure 3.**
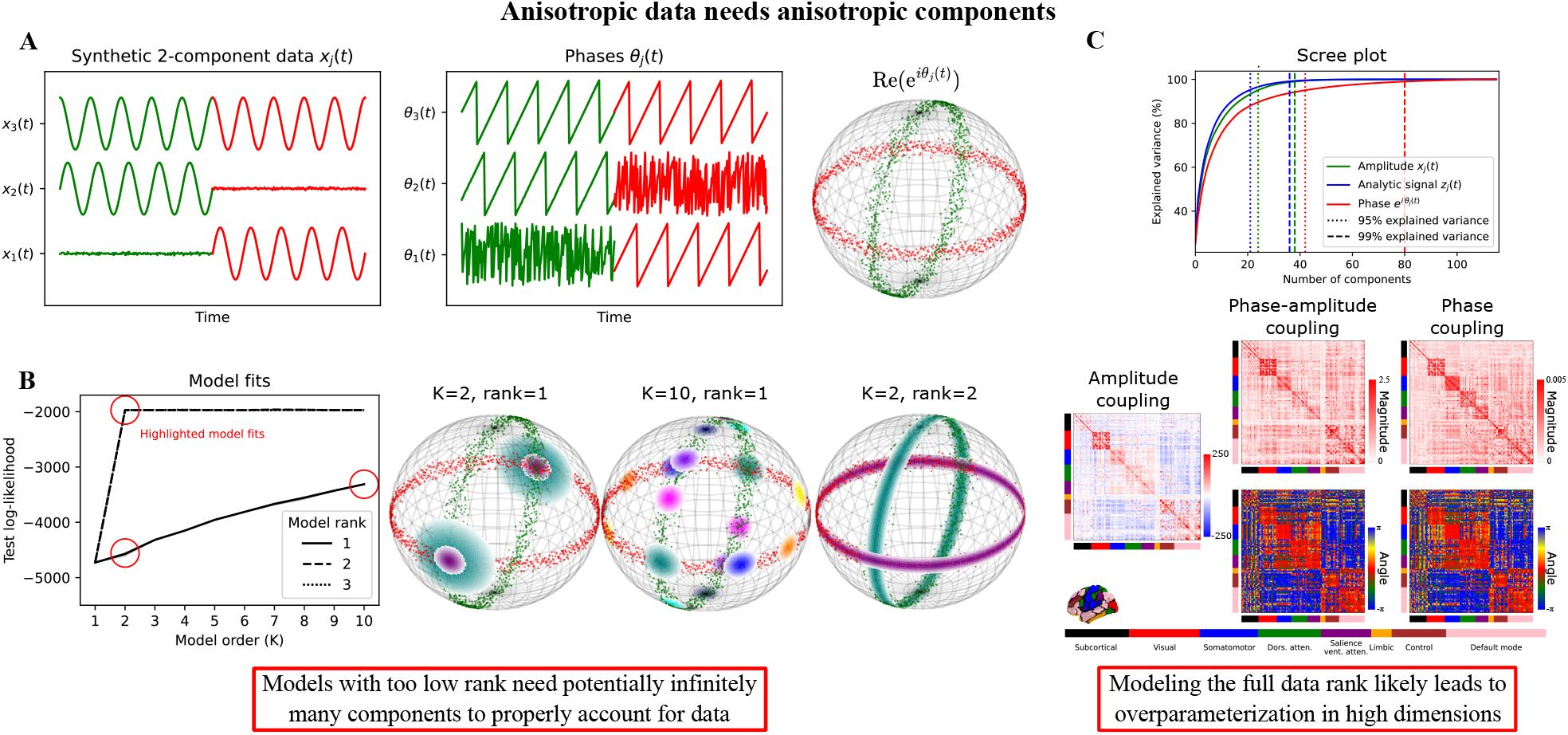
Non-unit rank models are needed to account for anisotropy present in functional neuroimaging data. **A**): Real-valued synthetic anisotropic phase vectors in two synthetic components. **B**): The (real-valued) angular central Gaussian mixture model with rank-1 components fails to account for the data anisotropy and instead fits isotropic components along the data manifold. With rank-2 or rank-3 covariance, the model adequately covers the spherical subspace on which the data is distributed. **C**): Functional neuroimaging data is naturally grouped into networks, which are inadequately described by rank-1 components but likely overparameterized by full-rank models. The solution involves modeling a certain predefined rank *r* in between 1 and *p*. Coupling matrices show global average covariances across 255 resting-state Human Connectome Project subjects for amplitude covariance from *x*(*t*), phase-amplitude coupling from *z*(*t*), and phase-only coupling from *u*(*t*).

To account for data anisotropy in models while avoiding overparameterization, the covariance matrix parameter **Ψ** may be reparameterized using a low-rank matrix plus the identity matrix, i.e., the so-called low-rank-plus-diagonal reparameterization: **Ψ** ≈ **Z** = **MM**^H^ + **I**. Here, **M** ∈ ℂ^*p*×*r*^ is a rank-*r* matrix, **I** is the identity matrix, and (·)^H^ is the complex conjugate transpose, reducing to the transpose for real-valued **M** (Olsen, Ortvald, et al. 2023). This allows explicitly controlling the complexity of connectivity captured by the components of the mixture model. For some distributions, such as the Gaussian distribution, a scale-parameter on the identity matrix is also required (Methods).

A mixture model with rank-controlled CACG distributions (real-valued for visualization purposes) as mixture components capture synthetic data to differing degrees (Fig. 3B): Rank-1 components are inherently isotropic, for which potentially infinitely many mixture components are needed to account for the anisotropy of the data. Instead, two rank-2 mixture components sufficiently explain the data. Additional components or increased component rank do not provide any gain in the predictive likelihood; such additional parameters are redundant and should be removed in keeping with the law of parsimony. Of note, many K-means models and matrix decompositions have rank-1 components and should thus be used with caution if the data is anisotropic.

Intercorrelations in fMRI data are seen by the naturally occurring network groupings of brain areas requiring modeling covariances beyond a rank-1 representation (Fig. 3C). Contrarily, a full covariance parameter **Ψ** can become singular leading to exploding normalization constants or modeling excessive numbers of parameters without explanatory gain.

To assess the practical implications of component ranks, we controlled the phase information in resting-state fMRI data from one subject by manipulating the phase in the Fourier domain into *K* = 5 segments (Supporting Fig. S2). Model performance was observed to stagnate around *r* = 10 with *p* = 116, indicating that unit-rank models underparameterize the data and full-rank models likely overparameterize. CACG mixtures outperformed all other models in recognizing the ground truth segments, showing their superiority in detecting phase changes in data unrelated to amplitude. Furthermore, we observed no performance benefits of using hidden Markov models over mixture models (Supporting Fig. S3) and therefore opted to only present mixture model results for the rest of the study. These mixtures were estimated using nonlinear optimization in PyTorch as we observed no benefit of analytical estimation via the expectation-maximization algorithm (Supporting Fig. S3).

### Phase coherence mixture modeling provides meaningful partitions into task-related functional states

Next, we investigated whether a low-rank phase coupling mixture model can extract meaningful task-related patterns from HCP fMRI (emotion, gambling, language, motor, relational, social, and working memory). Each task scan included both active and resting periods. While all blocks except the first pre-stimulus resting period were used for training the model, overlap with the true task label was only evaluated within task blocks corresponding to 1400 of 1870 total task volumes per subject. We applied the hold-out method and left 99 subjects out for testing (one subject excluded for not having all tasks completed) keeping 155 subjects for training. We fitted seven-component mixture models with varying component rank to the pooled training data (155 · 1870 fMRI volumes, no labels) and subsequently evaluated test-set posterior probability overlap with true task labels using normalized mutual information (NMI, Supporting Information) as well as the volume-wise and scan-wise prediction accuracy of the seven classes. Each model was run 10 times; only the best log-likelihood fit was analyzed (Fig. 4).

**Figure 4.**
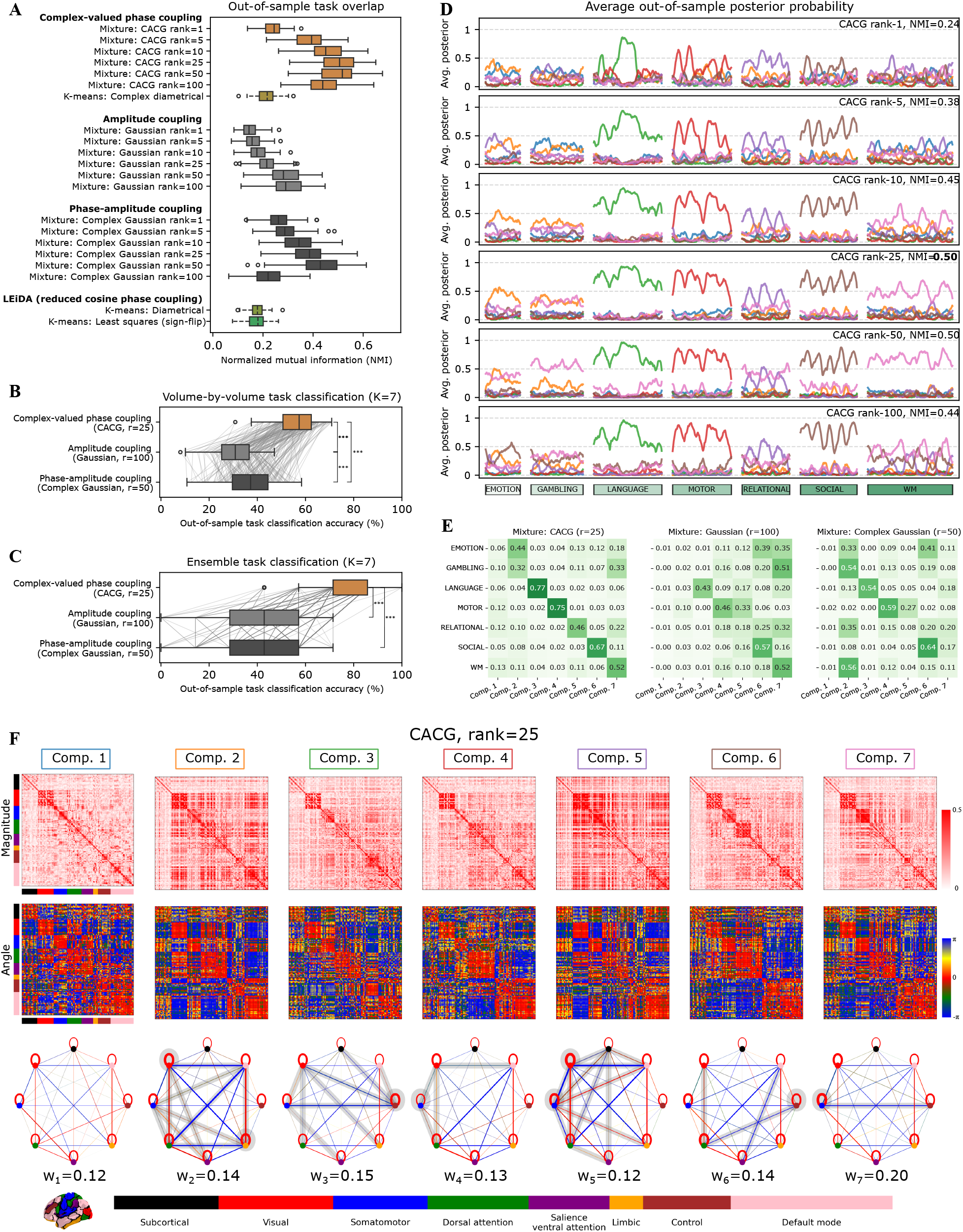
Seven-component mixture models trained on the seven task fMRI scans from the 155 subjects from the human connectome project (HCP) and evaluated on 99 held-out subjects. **A**): Overlap between test-set posterior probability and the true task labels measured using normalized mutual information. **B**): Volume-by-volume task classification accuracy using binarized test set posterior probabilities. Components were matched to tasks based on their average performance across train subjects. The average classification accuracy across volumes and scans is shown. **C**): Same as B), with only one predicted component per scan based on the average within-scan posterior. While models were trained on all volumes, NMI and accuracies were only evaluated within task blocks. In B) and C), ** *p <* 0.01, *** *p <* 0.001 indicates significance in a Wilcoxon signed rank test, Bonferroni-corrected for three comparisons. **D**): The average test posterior of each of the seven components (one color for each) of the CACG mixture model at different component ranks. The x-axis represents time with whitespace demarcating different task scans. Boxes indicate the name of the task and box widths are proportional to the number of volumes in the task scan. **E**): Test-set confusion matrices after ordering components to their most likely task as measured by train accuracy. **F**): The seven components learned using the best-performing model, i.e., the CACG mixture model with rank-25 components. In the network depiction, the color of edges corresponds to the average interregional “angle” and the line thickness to the average interregional magnitude in 8 canonical networks. Grey background for edges indicates that these are distinct in comparison with all other components (Hotelling’s T2-test, *p*_bonf_ *<* 0.05).

The best performing ranks were non-full (i.e., less than p) for all models: CACG for phase coupling (best rank r=25), zero-mean Gaussian for amplitude coupling (r=100, minor gain over r=50), and zero-mean complex Gaussian for phase-amplitude coupling (r=50) (Fig. 4A). The CACG (*r* = 25) achieved a test-NMI of 0.50 and volume-wise mean accuracy 55% (chance=14%), significantly outperforming the optimal Gaussian (36%) and complex Gaussian (37%) fits (Wilcoxon signed-rank *p*_bonf_ *<* 0.001). Scan-wise (one predicted label per scan) accuracy was higher (CACG: 75% significantly outperforming Gaussian: 41%, complex Gaussian: 43%, *p*_bonf_ *<* 0.001). All mixture models performed better than rank-1 K-means baselines.

To understand how model performance varied depending on the input data, we repeated the analysis at varying parcellation size and without global signal regression (GSR). CACG performance decreased without GSR (40% vs 55% with GSR), whereas amplitude models were less affected: 31% for Gaussian (*p*_bonf_ *<* 0.001 vs CACG) and 37% for complex Gaussian (*p*_bonf_ = 0.25 vs CACG, Supporting Fig. S4). At higher parcellation size, low-rank models performed similarly (rank-25 CACG accuracy at 232 regions: 55%) but high-rank models performed worse, indicating that controlling the number of estimated parameters becomes increasingly important with higher dimensionality. CACG was sensitive to an alternative denoising with temporal ICA, which reduced CACG accuracy to 47% while Gaussian (36%) and complex Gaussian (44%) were improved, suggesting GSR and preprocessing influence recoverable phase structure. Results were highly stable across dataset partitions (Supporting Fig. S5).

We visualized the average test set posterior and confusion matrix to examine how well each task was recovered by a single mixture component (Fig. 4D). The average test set posterior indicated that the language, motor, and social tasks were most recoverable, while emotion, relational, and working memory tasks were less discernible by the model; gambling was the hardest to detect, likely due to the very short event design (1s blocks) even leading this task to be excluded in other task classification studies on HCP data (Zhang et al. 2021; Rastegarnia et al. 2023). One mixture component (Component 1) behaved like a background state, indicating the presence of a non-task-related background component with spurious activation. The posterior also indicated consistent within-task fluctuations, e.g., five peaks for the social task, which is composed of five task blocks, and three peaks for the relational task, which is composed of three blocks. Confusions in CACG were mainly driven by gambling, relational, and working memory (Fig. 4E). The posteriors for the Gaussian and complex Gaussian showed lower task discernibility (Supporting Figs S7-8); these models were skewed in favor of a subset of components with high weight, likely caused by transient high-amplitude data variations becoming their own component, even if these are of low quantity.

Each component was characterized by a covariance matrix which we describe in terms of 8 canonical networks (Fig. 4F)(Thomas Yeo et al. 2011). All components were well-mixed across subjects (component weights ranged between 0.12–0.18). Few between-network connections were consistent across all components: the salience network was always coherent with somatomotor and dorsal attention networks, and default mode with limbic and control. The default network was always anticoherent with the dorsal attention network, visual with limbic, somatomotor with control, and salience with limbic. We summarize each CACG component by network connections that were significantly characteristic of that component versus all other components in terms of subject-wise connection strength (max p-value across component pairs using paired Hotelling’s T2-test (Hotelling 1931) on test subjects, Bonferroni corrected). The subcortical network only had two significant connections across components while the visual network had 17, the remaining networks had between 7-12 significant connections indicating neocortex-wide task engagement.

Component 1 (low task engagement, blue) showed generally low overall coupling with the notable exception of within-network visual coherence likely indicating the eyes-open condition. No connection in component 1 was significantly different from the same connection in other components. Component 2 (emotion, yellow) reflected a structured, arousal-like brain-wide state with broadly increased connection strength: limbic and visual networks displayed stronger within-network coherence and visual-dorsal attention, visual-salience, somatomotor-salience, and limbic-default coherences were increased; similarly with anticoherences: dorsal attention-default, dorsal attention-limbic, visual-default, visual-salience, somatomotor-dorsal attention, and somatomotor-default. Component 3 (language, green) was characterized by increased control-network coherence internally and pronounced anticoherence between control and subcortical, visual, and somatomotor networks. Unlike most components, visual and somatomotor networks were coherent, visual–dorsal attention showed anticoherence, whereas the somatomotor–salience coherence was reduced. The control network includes some language areas (left inferior frontal gyrus, angular and supramarginal gyri), so activation of this network is not surprising. Component 4 (motor, red) exhibited significantly reduced somatomotor within-network coherence and weakened anticoherence with visual regions; dorsal attention was coherent with somatomotor and consistently phase-shifted relative to visual, while visual was phase-shifted relative to default (a pattern unique among components). This finding suggests that area-specific motor engagement does not lead to network-wide somatomotor coherence. Component 5 (relational, purple) was marked by strengthened anticoherence between visual–default, visual–subcortical, visual–somatomotor, subcortical-dorsal attention, and somatomotor–dorsal attention, alongside coherent visual–dorsal attention, somatomotor-limbic, and somatomotor–default coupling (distinct from other components); portions of somatomotor–subcortical interactions were consistently phase-shifted. This component could indicate a perceptual integration mode, wherein visual information is mapped onto higher-order representations. Component 6 (social, brown) showed significantly increased control within-network coherence and enhanced visual–dorsal attention coherence, while control–dorsal attention relationships were anticoherent. The control-visual-attention coupling could indicate a structured reading of social gestures in the task. Finally, Component 7 (working memory, pink) was defined by within-network coherence in the somatomotor network together with anticoherence between somatomotor and control networks, indicating that the control network collaborates with the somatomotor network to actively memorize information.

Taken together, the CACG low-rank mixture model partitions the data into components that align with task structure and reveal uniquely interpretable changes in network coherence strength and phase polarity.

### Resting-state coherence is best described by a small set of consistent network patterns

On resting-state data, we trained models on the same 155 subjects and evaluated results on 100 test subjects. Specifically, we examined the number of mixture model components and component rank using three subject-specific metrics: 1) predictive log-likelihood indicating how well the test data matched the learned mixture distribution, 2) entropy of the average posterior indicating how “mixed” the components were for each subject, and 3) computational reproducibility indicating how sensitive the estimate was to changes in initialization influenced by the presence of local minima.

For the CACG mixture model, increasing the number of components altered likelihoods less than increasing component rank (Fig. 5A). When model rank tended towards the data’s dimensionality (*p* = 116), fewer components were supported by the test-set resting state data, and three components with rank-100 components achieved the highest predictive likelihood. These results suggest that resting-state brain fMRI data is not easily split into a high number of temporal segments that are representative across subjects and scans. Within-subject component mixing was slightly lower for ranks 50 and 100, indicating that the model tends to segregate subjects rather than individual volumes when components are of high rank. Computational reproducibility was highest for *K* ≤ 4 across all component ranks (*NMI >* 0.8 for most models). For higher model orders, estimates tended to fluctuate more, indicating that these models benefit from repeated initialization. Conclusions were similar for the Gaussian and complex Gaussian mixture models (Supporting Figs S8-9).

**Figure 5.**
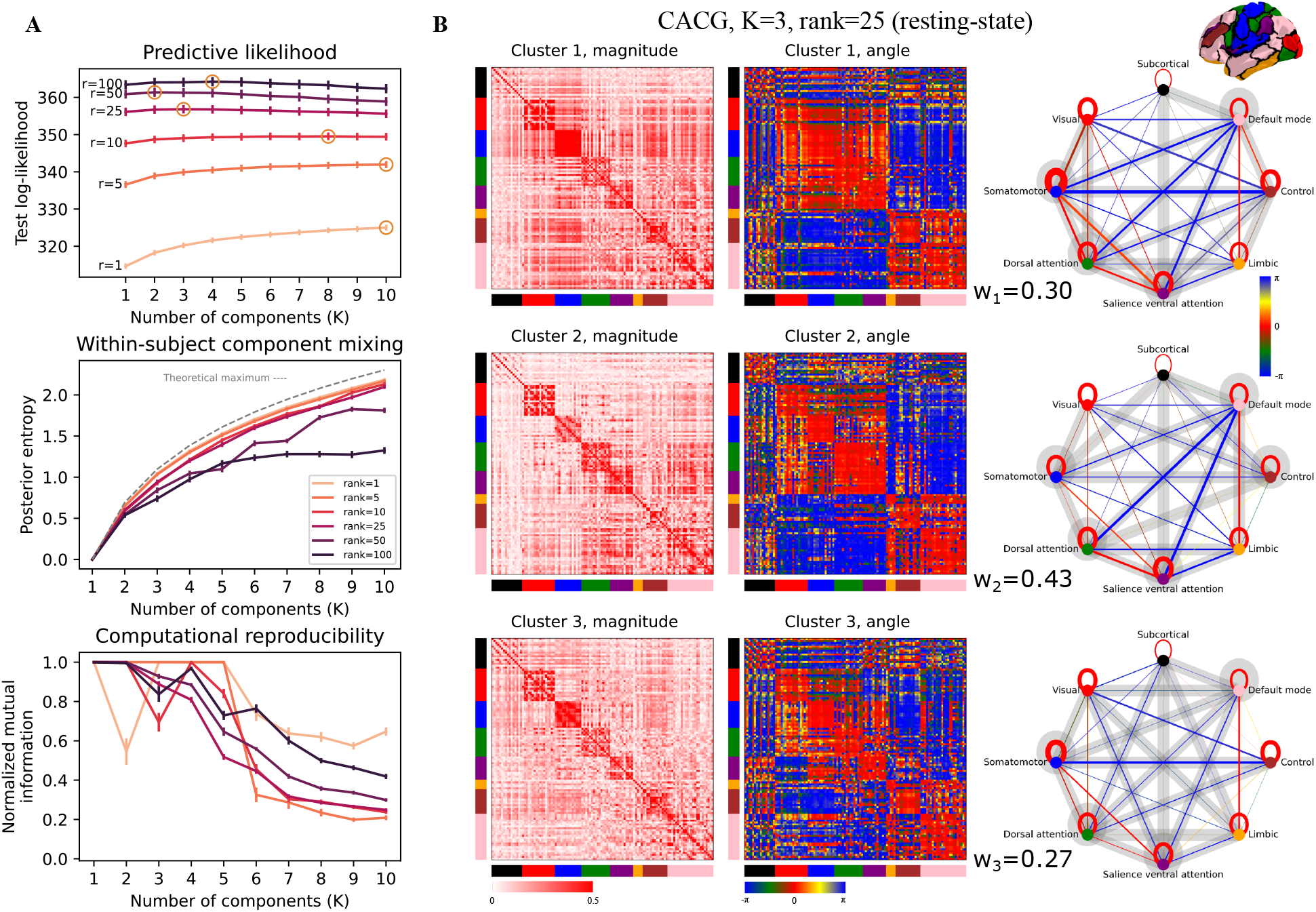
Mixture model parameter search and estimate on resting-state fMRI for the CACG model. **A**): Component rank and model order compared with out-of-sample test log-likelihood (normalized by the number of samples), posterior entropy indicating component mixing for each subject, and computational reproducibility upon repeated initializations of the same model; bars represent standard error across N=99 test subjects. **B**): CACG mixture estimate with K=3 and component ranks *r* = 25. In the network representation, average inter-network angle is represented by color, and magnitude by edge thickness. Significant connections (Hotelling’s T2-test, *p*_bonf_ *<* 0.05) are highlighted with a grey background.

We present the three-component CACG solution with rank-25 components, which had high within-subject mixing and computational reproducibility, in Fig. 5B. All components largely grouped sensorimotor and attention networks together against default-limbic and default-control networks. Given the fewer components, more connections were significant across components than for the task analysis; here we only describe selected connections. Component 1 was characterized by strengthened dorsal attention and somatomotor within- and between-network coherence and anticoherence between somatomotor and control networks, indicating task-positive activation potentially related to movements. Component 2, which was the most prominent, was characterized by decreased somatomotor/attention and control connectivity but otherwise displayed the conventional dissociation between default and task-positive networks. Component 3 was characterized by reduced dorsal attention and salience connectivity. The estimated components were similar to those estimated using Gaussian and complex Gaussian components (Supporting Figs. S8-9) though these seemingly had a wider distribution of network connectivity strengths.

GSR significantly altered the average resting-state global phase coherence map, revealing inter-regional anti-coherence in the angular representation (Supporting Fig. S11). Without GSR, the vast majority of the brain was, on average in this data, coherent. Following GSR, the average global phase coherence was predominantly close to zero (in-phase) or *π* (anti-phase). Visually, a CACG model with rank 10 was able to fit the global average phase coherence map well, with a visually negligible effect of increasing the rank. In contrast, a unit-rank model provided a notably simplified estimate of the global coherence map, essentially dividing the cortical areas into two chunks, suggesting that rank-1 models are too restrictive.

## Discussion

Here we explored and advanced multivariate probabilistic mixture modeling approaches for analyzing large-scale networks of phase coherence in their entirety, emphasizing the importance of discarding as little potentially valuable information as possible when modeling the data. Based on our results, we argue that simplifying data and models, such as by discarding the imaginary component of phase coherence, relying on leading eigenvectors, or utilizing rank-1 components, can result in loss of cognitively relevant information. Our findings show that appropriate statistical distributions that focus on phase coherence rather than signal amplitude better retrieve task information and provide a clean representation of brain dynamics. Importantly, our models were trained on label-free fMRI data and evaluated on unseen data from multiple subjects, indicating that the estimated models provide generalizable representations of dynamic brain connectivity. As such, our model may readily be used in future label-free studies to uncover, e.g., substages of sleep or common phase coherence patterns in people suffering from psychiatric disorders.

We introduced mixture models that were entirely agnostic to task labels, yet nonetheless supported accurate classification in held-out subjects (55% per-volume accuracy for the phase-synchronization model). Although task decoding was not a primary objective, this performance indicates that the learned unsupervised models capture cross-subject brain dynamics well enough to distinguish cognitively meaningful brain states. Prior work on task classification in the HCP dataset has relied largely on supervised or deep learning approaches. Classical supervised models have typically been applied to static functional connectomes (Hannum et al. 2023) or restricted to the immediate post-stimulus interval (Rastegarnia et al. 2023), limiting their ability to provide per-volume interpretability. Supervised discriminative methods also optimize directly for task labels and often depend on summary features, which further constrains mechanistic insight. Deep learning approaches achieve near-perfect decoding accuracy (Gupta, Lim, and Rajapakse 2022; X. Wang et al. 2019), but this is generally accompanied by reduced explainability of the underlying neural processes driving the predictions. In contrast, our framework offers a complementary, temporally resolved account of dynamic brain synchronization patterns. It preserves sample-level temporal resolution, provides meaningful task-related structure without requiring labels, and thereby bridges the gap between decoding performance and interpretability. By providing a principled way to extract and compare latent components across task and resting-state data, our approach enables future investigations to test specific hypotheses about behavioral relevance, physiological correlates, and underlying neural mechanisms. Applying this framework in conjunction with experimental perturbations or richer behavioral measures represents a natural next step.

On resting-state data, the CACG model with rank-25 parameter matrices achieved the highest performance using *K* = 3 clusters. Converging evidence from different models suggests that three networks are often used to describe brain connectivity, indicating a potentially parsimonious basis for resting-state brain function (Menon 2011; Bolt et al. 2022).

Through synthetic data, we illustrated that focusing solely on cosine phase coherence may mask valuable information about phase shifts between signals, a drawback remedied by complex-valued components (Fig. 2). The estimated seven-component CACG model for task fMRI data also suggested some valuable information in the imaginary component (green/yellow edges in Fig. 4F, “angle”). In the synthetic, phase-controlled and task-fMRI experiments, complex-valued models performed better than LEiDA, which models the leading eigenvector of the cosine phase coherence matrix and has been used extensively in the literature (Figs. 4, S1-S2, S4-S6). Taken together, constraining phase coherence models to the real domain risks discarding potential insights embedded in the imaginary component without obvious benefit.

In multivariate statistical modeling, attention to the geometry of the represented data is crucial. Eigenvectors, which are by definition unit norm and sign-invariant, are naturally associated with the projective, sign-symmetric unit hypersphere. Consequently, methods that treat such objects as unconstrained Euclidean vectors, such as least-squares or cosine-distance approaches, introduce geometric approximations, even with eigenvector sign-flipping as applied in LEiDA, which imposes non-affine restrictions on the data space (Olsen, Lykkebo-Valløe, et al. 2022). Some studies have suggested a K-means approach using either the *ℓ*_*l*_-norm, cosine distance, or Euclidean distance between the upper-triangular part of the full cosine phase coherence maps and centroids of the same size (Castro et al. 2024; Alteriis et al. 2024; Demertzi et al. 2019). While such approaches may be useful in practice, they do not explicitly model the manifold structure induced by phase data. Furthermore, representing only the triangular part of a symmetric matrix disregards its Riemannian structure leading to systematic modeling approximations. Models that more closely align with the relevant geometry of the data, and allow for components of variable rank, therefore provide a more suitable framework.

We estimated our models using both analytically derived update rules via expectation maximization (EM) as well as numerical optimization in PyTorch. We generally found the two methods to perform similarly (Supporting Fig. S3) with the latter being faster for large data sets. However, since the EM algorithm is more consistent (guarantees monotone convergence in many cases), we supply update rules and suggestions for model optimization for the CACG distribution based on EM, including low-rank-plus-diagonal reparameterization, in the Supporting Information. While our analyses were limited to low-dimensional data, low-rank-plus-diagonal reparameterization combined with gradual initialization of high-rank models (Methods) support the feasibility of future models in high, potentially native, resolution.

We showed that the global average phase coherence matrix without GSR is highly homogeneous (Supporting Fig. S10). Clustering cosine phase coherence maps without GSR typically results in one dominant global component with a high weight (Lord et al. 2019; Olsen, Lykkebo-Valløe, et al. 2022), while other clusters, previously referred to as ghost attractors (Vohryzek et al. 2020), carry less weight. The CACG distribution for phase data was markedly more affected by the removal of GSR than amplitude and phase-amplitude models (Supporting Fig. S4). Without GSR, fMRI data is often dominated by the global signal leading data to conform largely to this rank-1 component. Without access to amplitude information, the mixture of CACG distributions struggles more to discern subcomponents within such data. This is supported by high-rank CACG models, specifically, performing worse without GSR. In Gaussian and complex Gaussian models, amplitude variation allows for differentiating samples conforming to the global signal. Thus, for phase data, GSR is not just a denoising step to remove potential confounds, it is a necessity to extract meaningful phase coherence components. We believe that applying GSR, which removes more trivial coherence structures, will directly lead to learned components having more contrast, characterizing more detailed and relevant connectivity information.

A disadvantage of synchronization methods employing the Hilbert transform is the requirement of data to be narrowband for phase estimation. We used a filter in the range [0.03, 0.07]*Hz* based on previous studies (Glerean et al. 2012; Honari, Choe, and Lindquist 2021; ML et al. 2020; Pedersen et al. 2018), which is a relatively small restriction on the [0.01, 0.1]*Hz* range that the BOLD signal is generally accepted to fall within. Future studies should further investigate the effect of bandwidth on phase estimation in BOLD fMRI data, e.g., focusing on detecting phase slips (Pikovsky, Rosenblum, and Kurths 2001). Previously, the empirical mode decomposition has been suggested as a data-driven method to extract narrowband signal “modes” prior to the Hilbert transform (Honari and Lindquist 2022). However, it is difficult to reconcile frequency bands of interest across subjects and in practice, this method subdivides the frequency axis log-linearly with little indication of the relative relevance of each derived frequency band.

Besides the frequency range, our study has other limitations. The computational training speed increased markedly from K-means (time scale of seconds for 155 subjects) to mixture modeling (minutes) and hidden Markov modeling (hours). However, we did not exploit GPU-processing, which is readily available in PyTorch and could be implemented in future studies. We compared methods for phase coherence with zero-mean Gaussian mixture models for “raw” time-series data, however, in the literature, it is more common to see hidden Markov models for fMRI data, perhaps augmented with autoregressive parameters (Ahrends et al. 2022; Liégeois, Li, et al. 2019; Stevner et al. 2019; Vidaurre, Smith, and Woolrich 2017; Vidaurre, Quinn, et al. 2016). Here we observed little difference between mixture and hidden Markov modeling (Supporting Fig. S3). With the PyTorch stochastic estimation framework, future studies could very easily add an autoregressive emission specification to the presented models. We argue that phase coherence modeling is cleaner and more informative than amplitude models due to their insensitivity to outliers; however, future improved denoising models may narrow this gap. Future studies may also pursue a more in-depth analysis of phase-amplitude coupling, which is a large field of research in electrophysiology. Finally, we only examined one dataset; the estimated components may very well differ in different data.

## Conclusion

Modeling time-varying spatiotemporal coherence of functional brain imaging signals can provide meaningful insights about neural systems underlying cognition. Here we demonstrated that dynamic macroscale functional brain phase coherence networks can be modeled in their entirety considering mixture modeling approaches with 1) complex-valued and 2) non-unit rank centroids. We demonstrated the efficacy of the CACG mixture model on both synthetic and real resting-state and task fMRI. We suggest that future studies within phase coherence, whether in neuroscience or other fields, consider models such as the CACG distribution that account for anisotropy and phase coherence in its entirety. Our Phase Coherence Mixture Modeling (PCMM) toolbox is readily available for general-purpose phase coherence modeling: github.com/anders-s-olsen/PCMM.

## Methods

### Experimental data and preprocessing

We used fMRI data from 255 unrelated healthy subjects from the openly available Human Connectome Project (Elam et al. 2021; D. C. Van Essen et al. 2012). Ethics statements and imaging protocols may be found elsewhere (Barch et al. 2013; Smith et al. 2013); we downloaded the minimally preprocessed, multimodal surface matched task and resting-state data, the latter had also been ICA-FIX denoised (Glasser, Sotiropoulos, et al. 2013). We only used right-left phase encoding scans. Of the 255 subjects, 155 randomly selected individuals were used for training, while the remaining 100 subjects were exclusively used as an out-of-sample test set. For the 155 subjects, only one of two scans (REST1 or REST2, 1200 samples each, randomly selected) were included in the training set. One subject was discarded from the task analysis out-of-sample test set due to not having all task scans complete. We also evaluated a 2025-version of the HCP dataset using updated temporal and spatial ICA artifact classification schemes (Glasser, Coalson, et al. 2018). The task scans consisted of emotion (176 volumes), gambling (253), language (316), motor (284), relational (232), social (274), and working memory (405) tasks, see (Barch et al. 2013) for a full description of tasks. In each task scan we removed the initial part of the scan up until the first stimulus, corresponding to 15 volumes for emotion and 11 volumes for each of gambling, motor, relational, social, and working memory, such that the total number of volumes per subject across tasks was 1870.

The data was linearly detrended voxel-wise and subsequently spatially downsampled to 100 cortical and 16 subcortical regions using the Schaefer and Tian atlases (Schaefer et al. 2018; Tian et al. 2020). Subsequently, we bandpass-filtered the data in the range [0.03, 0.07]*Hz* with a 2nd-order zero-phase butterworth filter. Due to the relatively short acquisition lengths and long impulse response of the filter giving rise to edge effects from both filtering and the Hilbert transform, we prepended and appended each signal with 100 flipped samples of the signal itself, which corresponds exactly to 1% of the peak of the filter impulse response for TR=0.72s. The added samples were discarded after the Hilbert transform. Importantly, phase estimation via the Hilbert transform requires narrowband data; however, no direct guidelines for the required signal bandwidth exist, only rules of thumb such as requiring the bandwidth to be smaller than the mid-frequency (Feldman 2011) or twice the mid-frequency (Sun and Small 2009), both of which we enforce using 0.03−0.07*Hz* as our frequency range.

### Dynamic interregional phase coherence

Whereas the Pearson correlation coefficient between regional neuroimaging time-series implicitly assumes that the brain covariance matrix is static, time-varying phase coherence can be estimated via the Hilbert transform. Evaluating dynamic brain coherence implicitly assumes that the signal in all brain areas is of the same scale and that coactivity between regional time courses can be wholly represented by the coherence between their phases.

The regional neuroimaging signal *x*_*j*_(*t*) for region *j* ∈ {1, …, *p*} is extended into the complex domain using the Hilbert transform 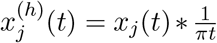, where ∗ is the convolution operator, to form the analytic signal

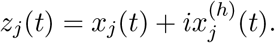

The Hilbert transform represents a phase shift by *π/*2 radians to every frequency in the signal. While the original signal *x*_*j*_(*t*) is oscillating along the time axis, the analytic signal *z*_*j*_(*t*) is revolving around the unit circle in the complex plane (Figs. 1-2). In the complex plane every point may be described using polar coordinates, i.e., as a combination of an angle to the positive real axis

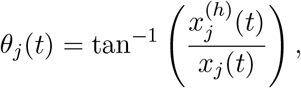

also called the instantaneous phase, and a distance from the origin 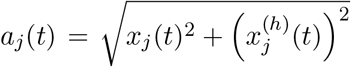 known as the instantaneous magnitude. The phase captures solely the oscillatory information in the signal while signal strength, including potential artifactual spikes, forms part of the signal amplitude. Whereas the Hilbert transform itself does not impose frequency restrictions on the signal, reliable estimation of phase requires that the signal be sufficiently narrowband (Boashash 1992). Furthermore, while the naming convention “instantaneous” implies that phases can be calculated using information from only a single time point, in reality the Hilbert transform suffers from the same estimation requirements for a sufficient number of samples and linearity/stationarity assumptions as the Fourier transform.

Each sample of the analytic signal is a complex number 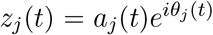, where the radius is ignored when exclusively considering phase coherence, i.e., *a*_*j*_(*t*) = 1. The phase *θ* and difference between two phases *θ*_1_ − *θ*_2_ are circular variables with modus 2*π*. The rectangular representation can therefore be used to assess coherence via *e*^*i*(*θ*1−*θ*2)^ and more generally, all pairwise complex instantaneous phase coherences can be represented by the matrix **A**_*t*_ with elements

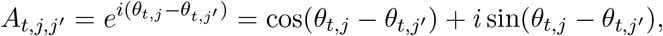

such that every element for nodes *j, j*^′^ ∈ {1, …, *p*} provides an estimate of the pairwise instantaneous synchrony between regions. Each element of **A**_*t*_ can be factored as 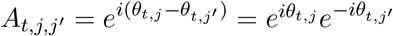, and thus, the matrix can be written as 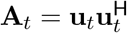, where 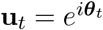 and (·)^H^ represents the complex conjugate transpose. Thus, the rank of **A**_*t*_ is 1, and instantaneous interregional phase coherence can be represented using a vector, **u**_*t*_, instead of a matrix with ^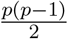^ unique elements, greatly simplifying the modeling of all pairwise coherences. Since diag(**A**_*t*_) = **1**, i.e., a vector of all ones, the only non-zero eigenvalue of **A**_*t*_ is *λ* = *p*. Thus, **u**_*t*_ may readily be rescaled to unit norm, 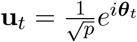. Combined with the fact that flipping the sign of **u**_*t*_ does not change **A**_*t*_, this means that **u**_*t*_ is an eigenvector and models of interregional phase coherence should therefore also be sign-invariant. Consequently, all pairwise instantaneous phases can be compactly represented using a complex-valued sign-invariant vector residing on the unit sphere, also known as the complex projective hyperplane ℂℙ^*p*−1^ = {**u**|**u** ∈ ℂ^*p*^, **u**^H^**u** = 1, −**u** = **u**}. Of note, phase vectors occupy a hypertorus embedded in the complex projective hyperplane. However, flexible likelihood-based models on hypertori require normalizing constants that are unavailable in closed form and computationally intractable in high dimensions. Therefore, we employ models for the complex projective hyperplane.

It is also possible to model phase coherence in the real domain by discarding the imaginary part of the complex phase coherence. Especially in fMRI research, the cosine phase coherence matrix has been widely studied, particularly in the context of the Leading Eigenvector Dynamics Analysis

(LEiDA) (Cabral et al. 2017) considering the real symmetric matrix 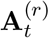 with elements

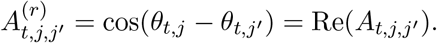

Notably, a matrix of cosine differences is rank-2 due to the angle difference identity

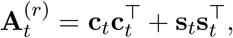

where **c**_*t*_ = cos ***θ***_*t*_ = Re(**u**_*t*_) and **s**_*t*_ = sin ***θ***_*t*_ = Im(**u**_*t*_), which we have shown previously in (Olsen, Lykkebo-Valløe, et al. 2022). Consequently, 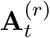 can be expressed in terms of two eigenvectors with non-zero eigenvalue. The leading eigenvector of the cosine phase coherence map automatically accounts for at least 50% of the variance and empirically captures anywhere between 50% and 100% (Alteriis et al. 2024). Several previous studies have focused on the leading eigenvector **u**_*t*,1_ of 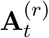 in the LEiDA-framework (Cabral et al. 2017). These eigenvectors have traditionally been sign-flipped, such that the majority of elements are of the same sign, and subsequently clustered using least-squares K-means (Cabral et al. 2017; Figueroa et al. 2019; Lord et al. 2019; Vohryzek et al. 2020). This approach is a heuristic, which we previously have shown may lead to inaccurate results (see Fig. S1 in (Olsen, Lykkebo-Valløe, et al. 2022)). In our previous work, we therefore suggested using diametrical clustering as a projective hyperplane alternative for LEiDA (Olsen, Lykkebo-Valløe, et al. 2022).

### Statistical distributions and low-rank reparameterization

Below we present statistical distributions for analyzing multivariate brain data from amplitude, phase, and phase-amplitude perspectives; the presented methods and their characteristics are summarized in Supporting Table S1. Further information on the CACG distribution may be found in the Supporting Information.

### Complex angular central Gaussian distribution

The complex angular central Gaussian (CACG) distribution for data on the complex projective hyperplane has density (Kent 1997; Tyler 1987)

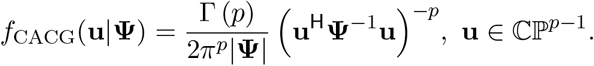

Here, **Ψ** ∈ ℂ^*p*×*p*^ is a Hermitian-symmetric positive definite matrix identifiable up to a positive scale factor (**Ψ** is scaled such that tr(**Ψ**) = *p* for convenience) and | · | is the matrix determinant. The CACG distribution is named based on its sampling procedure: samples from a centered multivariate complex normal distribution normalized to the sphere follows the CACG, i.e.,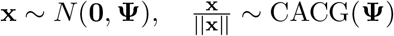 (Mardia and Jupp 1999). The Bingham distribution, which serves a similar purpose, includes a more complicated normalization constant (the confluent hypergeometric function of matrix argument), which becomes intractable for non-trivial dimensionality, rendering the CACG distribution an attractive alternative. Maximum likelihood estimation is deferred to the Supporting Information.

### Gaussian distribution

The real-valued zero-mean multivariate Gaussian distribution has the density

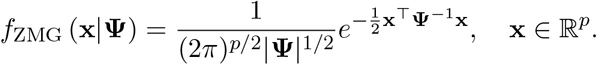

Similarly, the zero-mean complex-valued multivariate Gaussian distribution has the density

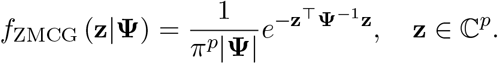

### Low-rank-plus-diagonal reparameterization of Ψ

Estimating covariance matrix parameters can be challenging due to the near-quadratic increase in parameters with dimensionality. Moreover, high-resolution brain connectivity networks exhibit rank deficiency due to related areas having highly correlated time-series. To address this, a low-rank-plus-diagonal reparameterization **Ψ** ≈ *γ* **(MM**^H^ + **I)**, with **M** ∈ ℂ^*p*×*r*^ (or **M** ∈ ℝ^*p*×*r*^ for the Gaussian), allows capturing a pre-specified rank *r* ≤ *p* of the learned parameter matrix (Olsen, Ortvald, et al. 2023). The constant *γ* ∈ ℝ for the CACG distribution corresponds to a trace-normalization, i.e., 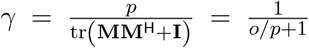 where 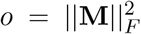, which does not affect mixture model results due to the scale-invariance of the CACG distribution. For the zero-mean Gaussian and complex Gaussian distributions, that are not scale-invariant, we used the reparameterization **Ψ** ≈ **MM**^H^ + *γ***I** such that the scale parameter affects the relationship between the added identity matrix and **M**. The use of low-rank-plus-diagonal reparameterization required a re-derivation of existing analytical maximization updates to be with respect to **M** as detailed in the Supporting Information.

### Probabilistic mixture modeling

Mixture modeling involves computing parameters for *K* components, each corresponding to a unique probability distribution mixed with component weights 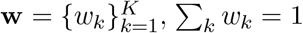. The mixture den-sity for each observation **y**_*t*_, which may correspond to, e.g., a (complex) eigenvector **y**_*t*_ = **u**_*t*_ (phase coupling, CACG), a scaled phase vector **y**_*t*_ = **z**_*t*_ (phase-amplitude coupling, complex Gaussian), the time-series **y**_*t*_ = **x**_*t*_ (amplitude coupling, Gaussian), is defined as

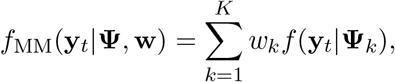

where **Ψ**_*k*_ is the covariance matrix of the *k*^*th*^ component.

We utilized two estimation schemes for mixture models involving these distributions: expectation-maximization (EM) and direct likelihood optimization. Both methods optimize the mixture model log-likelihood function evaluated over the entire data set 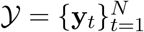

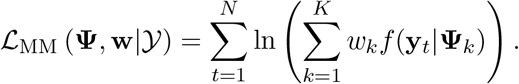

EM computes assignment probabilities (i.e., state responsibilities a.k.a. posterior) *β*_*kt*_ for sample *t* and component *k* in the expectation (E-step) according to

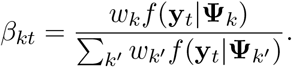

These probabilities are used during the maximization (M-step) to guide analytical parameter updates. In contrast, the direct optimization procedure only evaluates the log-likelihood function and updates parameters directly using an optimizer and automatic differentiation. Although the E-step is not required for direct numerical optimization, it is still used to compute posterior probabilities based on the estimated parameters.

### Implementation details

We implemented mixture models with analytical maximization in NumPy and numerical optimization in PyTorch, where the latter also included the possibility for computing hidden Markov models (Supporting Information). Recent advances in fast numerical gradient optimization frameworks including automatic differentiation in PyTorch has enabled fast non-linear model inference while only requiring the specification of an objective function, in this case the log-likelihood function. An advantage of direct numerical optimization when compared to the more conventional EM inference procedure is that the CACG and Gaussian models with low-rank-plus-diagonal reparameterization all require iterative fixed-point estimation for analytical maximization in the M-step. Conversely, direct numerical optimization requires choosing a learning rate and learning algorithm; Here we used the ADAM optimizer (Kingma and Ba 2014) with LR = 0.1 in correspondence with adjacent studies (Olsen, Ortvald, et al. 2023; Olsen, J. D. Nielsen, and Mørup 2024).

Parameter constraints for numerical optimization were implemented through reparametrization of unconstrained variables to comply with the imposed constraints: We used a softmax function for the mixture weights 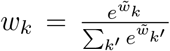 and a softplus function for the *γ*-parameter for Gaussian distributions to ensure non-negativity.

Mixture models, both using analytical and numerical optimization, were evaluated until a relative difference between the best and second-best iteration (divided by the best) in the latter 50 epochs fell below a tolerance level of tol = 10^−10^. While the EM algorithm typically guarantees monotone convergence, this is not the case for the low-rank-plus-diagonal reparameterizations, since these rely on iterative optimization procedures in the M-step. Analytical (EM) fixed-point updates for CACG and Gaussian distributions were evaluated until an absolute tolerance of 10^−10^.

### K-means equivalents

To provide a comparison to the probabilistic mixture models and to function as initialization to these, we utilized existing hard assignment K-means models for the projective hyperplane and introduced a complex-valued version of diametrical clustering (Dhillon, Marcotte, and Roshan 2003) for the complex projective hyperplane.

### Initialization schemes

Given the vast array of available models, gradually increasing model complexity may aid in ensuring model convergence to a suitable optimum. The K-means algorithms were initialized using their ++-versions, where an initial centroid is picked randomly and the remaining *K* − 1 centroids iteratively picked based on a probability distribution proportional to the distance to existing centroids, ensuring that the starting points were spread far apart (Arthur and Vassilvitskii 2007). This scheme provided slightly better results than initializing centroids via the random uniform distribution (Supporting Fig. S11). Probabilistic mixture models were initialized using centroids derived from K-means. The low-rank matrix **M** was initialized, given a partition of the data set **y**_*t*_ ∈ C_*k*_, e.g., from K-means, as

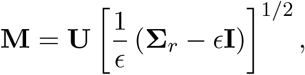

where **UΣ V**^H^ = **Y** is the singular value decomposition and **Y** = 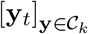 are the samples assigned to cluster *k* stacked horizontally. **Σ** _*r*_ is the diagonal matrix of singular values 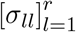, where all other values than the leading *r* singular values were set to zero, and 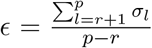, assuming the number of assigned samples to cluster *k* is *n*_*k*_ *> p*. Mixture component weights **w** were initialized based on the initialization partition. For the Gaussian distributions, *γ* was initialized to the same value as *ϵ*.

We also investigated a scenario where high-rank mixtures were initialized by their lower-rank counterparts. For instance, given the parameter **M**_*h*_ of a rank *h < r* model, the first *h* columns of **M** *h* could be initialized as **M**_*h*_ and the remaining *r*−*h* columns as the singular value decomposition on the*h* residual matrix after projecting out the existing columns, i.e., 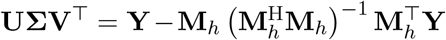 Gradual initialization provided improved results (Supporting Fig. S3).

### Model evaluation

We evaluated our methods on both synthetic, phase-manipulated, and real task and resting-state fMRI data. To compare posterior probabilities to a known ground truth, we used normalized mutual information (NMI), which is invariant to permutation in clustering labels (see Supporting Information for the definition of NMI).

Synthetic data was generated from two true three-dimensional components with consistent phase shifts and an added noise level *σ* = *π/*16, …, *π/*2, *n* = 1000 samples for each component (Supporting Fig. S1). Models were fitted for each noise level separately and NMI evaluated against the true component labels.

To evaluate performance upon increasing model complexity (Supporting Fig S2), we used phase-controlled 116-dimensional resting-state data from one subject and one scan, i.e., *N* = 1200 data points corresponding to 15 minutes of data acquisition. First we divided both *N* and *p* into *K* = 5 groups, such that cluster 1 corresponded to the first 240 time points and the first 23 regions, cluster 2 corresponded to time points 241 to 480 and regions 24-46 and so on. Then we computed, for all regions and temporal windows, the Fourier spectrum and if the region and window corresponded to one of the clusters, the phase information for the relevant frequencies was set to a pre-specified constant phase *α*_*j*_ ∈ [0; 2*π*]. If not, the phase information within the frequency band was randomized. We subsequently evaluated NMI between the posterior probability and one-hot encoded true labels.

Task fMRI scans were concatenated for train and test subjects into a full train and test set. Seven-component models were estimated on the full train set without access to labels, and subsequently the posterior probabilities were computed on the test set. Separate mixture models were estimated with component ranks *r* ∈ [1, 5, 10, 25, 50, 100]. Only the best of 10 estimates, evaluated using train set likelihood, were analyzed. We evaluated the overlap between true task labels and the posterior probability using NMI and classification accuracy only within task blocks defined by HCP, reducing the number of evaluated volumes across all task scans from 1870 to 1400 for each subject. To compute a classification accuracy, we binarized the posterior at each sample to obtain an estimated “class” prediction, which we compared with true task labels within task blocks, i.e., no additional classifier was trained. We first aligned each component with the task label that it corresponded to the most in the train set (average across all subjects) by solving a linear sum assignment problem and then evaluated classification accuracy on the test set. We also aggregated the posterior within each task scan to obtain an aggregated class prediction and computed a corresponding ensemble seven-class classification accuracy. Classification accuracies were tested between models using a Wilcoxon signed rank test Bonferroni-corrected for three comparisons.

Component covariance matrices were reduced from 116 parcels to 8 cortical and subcortical networks (Figs. 4F, S5-6). To assess which edges were specific to individual components, we first computed weighted complex-valued covariance matrices for each test subject and component using the test set posterior probabilities as weights. For each component, complex-valued covariance elements were compared to the corresponding elements in all other components using pairwise paired Hotelling’s T2-test (Hotelling 1931) on the test subjects (N=99 for task, N=100 for resting-state). For diagonal elements (self-connections), and the non-complex Gaussian model, we instead applied paired t-tests. This resulted in *p*(*p*−1)*/*2· *K*(*K* −1)*/*2 (*p* = 8 canonical networks) statistical tests per model for each of the three models considered (CACG, Gaussian, complex Gaussian). To summarize component specificity for a given network edge, we retained the maximum p-value across all *K* − 1 component pairs involving that edge. The combined *K* · *p*(*p*−1)*/*2 p-values were Bonferroni corrected for multiple comparisons.

On resting-state fMRI data we fitted models with varying model order *K* and component rank *r*. Each model was fitted 10 times and computational reproducibility evaluated using NMI between the posterior of all pairs of estimates. On the best estimate (evaluated using train set likelihood), we computed, for each test subject, the average sample log-likelihood and posterior entropy, computed 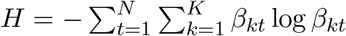, where *N* = 2400 is the number of volumes per test subject.

## Acknowledgements

This work was supported by the Danish Pioneer Centre for AI, DNRF grant number P1.

## Author contributions

ASO and MM conceived the study and developed the theoretical framework. ASO performed the experiments, wrote the toolbox, and drafted the manuscript. AB aided in model development and PMF aided in analysis of results. MM and PMF supervised the project. All authors contributed to interpretation of results and reviewed and approved the final version of the manuscript.

## Data availability

All processing and analysis scripts used in the present study, implemented primarily in Python, are made available at github.com/anders-s-olsen/PCMM. Human connectome project data are readily available online.

## References

Ahrends, C et al. (May 2022). “Data and model considerations for estimating time-varying functional connectivity in fMRI”. In: NeuroImage 252, p. 119026. ISSN: 1053-8119. doi: 10.1016/j.neuroimage.2022.119026. URL: https://www.sciencedirect.com/science/article/pii/S1053811922001550 (visited on 10/26/2024).

Allen, Elena A. et al. (Mar. 2014). “Tracking whole-brain connectivity dynamics in the resting state”. In: Cerebral Cortex 24.3. Publisher: Cereb Cortex, pp. 663–676. ISSN: 10473211. doi: 10.1093/cercor/bhs352. URL: https://pubmed.ncbi.nlm.nih.gov/23146964/ (visited on 07/27/2021).

Alteriis, Giuseppe de et al. (Mar. 2024). “EiDA: A lossless approach for dynamic functional connectivity; application to fMRI data of a model of ageing”. In: Imaging Neuroscience 2, pp. 1–22. ISSN: 2837-6056. doi: 10.1162/imag_a_00113. URL: https://doi.org/10.1162/imag_a_00113 (visited on 05/29/2024).

Arthur, David and Sergei Vassilvitskii (2007). “K-means++: The advantages of careful seeding”. In: Proceedings of the Annual ACM-SIAM Symposium on Discrete Algorithms. Vol. 07-09-Janu, pp. 1027–1035. ISBN: 978-0-89871-624-5. (Visited on 07/23/2021).

Barch, Deanna M. et al. (May 2013). “Function in the Human Connectome: Task-fMRI and Individual Differences in Behavior”. en. In: NeuroImage 80, p. 169. doi: 10.1016/j.neuroimage.2013.05.033. URL: https://pmc.ncbi.nlm.nih.gov/articles/PMC4011498/ (visited on 10/27/2024).

Bastos André M. and Jan Mathijs Schoffelen (Jan. 2016). “A tutorial review of functional connectivity analysis methods and their interpretational pitfalls”. In: Frontiers in Systems Neuroscience 9.JAN2016. Publisher: Front Syst Neurosci. ISSN: 16625137. doi: 10.3389/fnsys.2015.00175. URL: https://pubmed.ncbi.nlm.nih.gov/26778976/ (visited on 07/23/2021).

Biswal, Bharat et al. (1995). “Functional connectivity in the motor cortex of resting human brain using echo-planar mri”. In: Magnetic Resonance in Medicine 34.4. Publisher: Magn Reson Med, pp. 537–541. ISSN: 15222594. doi: 10.1002/mrm.1910340409. URL: https://pubmed.ncbi.nlm.nih.gov/8524021/ (visited on 09/08/2021).

Boashash, B. (Apr. 1992). “Estimating and interpreting the instantaneous frequency of a signal. I. Fundamentals”. In: Proceedings of the IEEE 80.4. Conference Name: Proceedings of the IEEE, pp. 520–538. ISSN: 1558-2256. doi: 10.1109/5.135376. URL: https://ieeexplore.ieee.org/document/135376 (visited on 01/10/2025).

Bola, Michal and Bernhard A. Sabel (July 2015). “Dynamic reorganization of brain functional networks during cognition”. In: NeuroImage 114, pp. 398–413. ISSN: 1053-8119. doi: 10.1016/j.neuroimage.2015.03.057. URL: https://www.sciencedirect.com/science/article/pii/S1053811915002530 (visited on 11/15/2024).

Bolt, Taylor et al. (2022). “A parsimonious description of global functional brain organization in three spatiotemporal patterns”. In: Nature Neuroscience. Publisher: Springer US. ISSN: 1097-6256. doi: 10.1038/s41593-022-01118-1. URL: https://www.nature.com/articles/s41593-022-01118-1.

Cabral, Joana et al. (2017). “Cognitive performance in healthy older adults relates to spontaneous switching between states of functional connectivity during rest”. In: Scientific Reports 7.1, pp. 1– 13. ISSN: 20452322. doi: 10.1038/s41598-017-05425-7.

Castro, Pablo et al. (Sept. 2024). “Dynamical structure-function correlations provide robust and generalizable signatures of consciousness in humans”. en. In: Communications Biology 7.1. Publisher: Nature Publishing Group, pp. 1–12. ISSN: 2399-3642. doi: 10.1038/s42003-024-06858-3. URL: https://www.nature.com/articles/s42003-024-06858-3 (visited on 10/30/2024).

Chen, Jingyuan E., Mikail Rubinov, and Catie Chang (Nov. 2017). “Methods and Considerations for Dynamic Analysis of Functional MR Imaging Data”. In: Neuroimaging Clinics of North America. Functional Connectivity 27.4, pp. 547–560. ISSN: 1052-5149. doi: 10.1016/j.nic.2017.06.009. URL: https://www.sciencedirect.com/science/article/pii/S1052514917300679 (visited on 11/15/2024).

Choe, Ann S. et al. (Sept. 2017). “Comparing test-retest reliability of dynamic functional connectivity methods”. In: NeuroImage 158, pp. 155–175. ISSN: 1053-8119. doi: 10.1016/j.neuroimage.2017.07.005. URL: https://www.sciencedirect.com/science/article/pii/S1053811917305736 (visited on 11/09/2024).

Coombes, Stephen and Paul C. Bressloff (2005). Bursting: The Genesis of Rhythm in the Nervous System. en. Google-Books-ID: OQN0xuOMKN0C. World Scientific. ISBN: 978-981-256-506-8.

Demertzi, A. et al. (Feb. 2019). “Human consciousness is supported by dynamic complex patterns of brain signal coordination”. In: Science Advances 5.2. Publisher: American Association for the Advancement of Science, eaat7603. doi: 10.1126/sciadv.aat7603. URL: https://www.science.org/doi/10.1126/sciadv.aat7603 (visited on 10/30/2024).

Dhillon, Inderjit S., Edward M. Marcotte, and Usman Roshan (2003). “Diametrical clustering for identifying anti-correlated gene clusters”. In: Bioinformatics 19.13, pp. 1612–1619. ISSN: 13674803. doi: 10.1093/bioinformatics/btg209.

Elam, Jennifer Stine et al. (Dec. 2021). “The Human Connectome Project: A retrospective”. In: NeuroImage 244, p. 118543. ISSN: 1053-8119. doi: 10.1016/j.neuroimage.2021.118543. URL: https://www.sciencedirect.com/science/article/pii/S1053811921008168 (visited on 05/15/2024).

Feldman, Michael (Apr. 2011). “Hilbert transform in vibration analysis”. In: Mechanical Systems and Signal Processing 25.3, pp. 735–802. ISSN: 0888-3270. doi: 10.1016/j.ymssp.2010.07.018. URL: https://www.sciencedirect.com/science/article/pii/S0888327010002542 (visited on 01/11/2025).

Figueroa, Caroline A. et al. (2019). “Altered ability to access a clinically relevant control network in patients remitted from major depressive disorder”. In: Human Brain Mapping 40.9, pp. 2771–2786. ISSN: 10970193. doi: 10.1002/hbm.24559.

Glasser, Matthew F., Timothy S. Coalson, et al. (Nov. 2018). “Using temporal ICA to selectively remove global noise while preserving global signal in functional MRI data”. In: NeuroImage 181, pp. 692–717. ISSN: 1053-8119. doi: 10.1016/j.neuroimage.2018.04.076. URL: https://www.sciencedirect.com/science/article/pii/S1053811918303963 (visited on 12/17/2025).

Glasser, Matthew F., Stamatios N. Sotiropoulos, et al. (Oct. 2013). “The minimal preprocessing pipelines for the Human Connectome Project”. In: NeuroImage. Mapping the Connectome 80, pp. 105–124. ISSN: 1053-8119. doi: 10.1016/j.neuroimage.2013.04.127. URL: https://www.sciencedirect.com/science/article/pii/S1053811913005053 (visited on 12/23/2023).

Glerean, Enrico et al. (Apr. 2012). “Functional Magnetic Resonance Imaging Phase Synchronization as a Measure of Dynamic Functional Connectivity”. In: Brain Connectivity 2.2, pp. 91–101. ISSN: 2158-0014. doi: 10.1089/brain.2011.0068. URL: https://www.ncbi.nlm.nih.gov/pmc/articles/PMC3624768/ (visited on 05/29/2024).

Gupta, Sukrit, Marcus Lim, and Jagath C. Rajapakse (2022). “Decoding task specific and task general functional architectures of the brain”. en. In: Human Brain Mapping 43.9. eprint: https://onlinelibrary.wiley.com/doi/pdf/10.1002/hbm.25817, pp. 2801–2816. ISSN: 1097-0193. doi: 10.1002/hbm.25817. URL: https://onlinelibrary.wiley.com/doi/abs/10.1002/hbm.25817 (visited on 12/08/2025).

Hannum, Andrew et al. (2023). “High-accuracy machine learning techniques for functional connectome fingerprinting and cognitive state decoding”. en. In: Human Brain Mapping 44.16. eprint: https://onlinelibrary.wiley.com/doi/pdf/10.1002/hbm.26423, pp. 5294–5308. ISSN: 1097-0193. doi: 10.1002/hbm.26423. URL: https://onlinelibrary.wiley.com/doi/abs/10.1002/hbm.26423 (visited on 12/08/2025).

Hindriks, R. et al. (Feb. 2016). “Can sliding-window correlations reveal dynamic functional connectivity in resting-state fMRI?” In: NeuroImage 127, pp. 242–256. ISSN: 1053-8119. doi: 10.1016/j.neuroimage.2015.11.055. URL: https://www.sciencedirect.com/science/article/pii/S1053811915010782 (visited on 10/30/2024).

Honari, Hamed, Ann S. Choe, and Martin A. Lindquist (Mar. 2021). “Evaluating phase synchronization methods in fMRI: A comparison study and new approaches”. In: NeuroImage 228, p. 117704. ISSN: 1053-8119. doi: 10.1016/j.neuroimage.2020.117704. URL: https://www.sciencedirect.com/science/article/pii/S1053811920311897 (visited on 10/13/2024).

Honari, Hamed and Martin A. Lindquist (Nov. 2022). “Mode decomposition-based time-varying phase synchronization for fMRI”. In: NeuroImage 261, p. 119519. ISSN: 1053-8119. doi: 10.1016/j.neuroimage.2022.119519. URL: https://www.sciencedirect.com/science/article/pii/S1053811922006346 (visited on 10/13/2024).

Hotelling, Harold (Aug. 1931). “The Generalization of Student’s Ratio”. In: The Annals of Mathematical Statistics 2.3. Publisher: Institute of Mathematical Statistics, pp. 360–378. ISSN: 0003-4851, 2168-8990. doi: 10.1214/aoms/1177732979. URL: https://projecteuclid.org/journals/annals-of-mathematical-statistics/volume-2/issue-3/The-Generalization-of-Students-Ratio/10.1214/aoms/1177732979.full (visited on 01/05/2026).

Kent, John T. (Feb. 1997). “Data analysis for shapes and images”. In: Journal of Statistical Planning and Inference. Robust Statistics and Data Analysis, Part II 57.2, pp. 181–193. ISSN: 0378-3758. doi: 10.1016/S0378-3758(96)00043-2. URL: https://www.sciencedirect.com/science/article/pii/S0378375896000432 (visited on 08/06/2024).

Kingma, Diederik P. and Jimmy Lei Ba (Dec. 2014). “Adam: A Method for Stochastic Optimization”. In: 3rd International Conference on Learning Representations, ICLR 2015 - Conference Track Proceedings. arXiv: 1412.6980 Publisher: International Conference on Learning Representations, ICLR. doi: 10.48550/arxiv.1412.6980. URL: https://arxiv.org/abs/1412.6980v9 (visited on 03/02/2023).

Kirschstein, Timo and Rüdiger Köhling (July 2009). “What is the Source of the EEG?” en. In: Clinical EEG and Neuroscience 40.3. Publisher: SAGE Publications Inc, pp. 146–149. ISSN: 1550-0594. doi: 10.1177/155005940904000305. URL: https://doi.org/10.1177/155005940904000305 (visited on 11/09/2024).

Laumann, Timothy O., Abraham Z. Snyder, and Caterina Gratton (Oct. 2024). “Challenges in the measurement and interpretation of dynamic functional connectivity”. In: Imaging Neuroscience. ISSN: 2837-6056. doi: 10.1162/imag_a_00366. URL: : https://doi.org/10.1162/imag_a_00366 (visited on 11/15/2024).

Laumann, Timothy O., Abraham Z. Snyder, Anish Mitra, et al. (Oct. 2017). “On the Stability of BOLD fMRI Correlations”. In: Cerebral Cortex 27.10, pp. 4719–4732. ISSN: 1047-3211. doi: 10.1093/cercor/bhw265. URL: https://doi.org/10.1093/cercor/bhw265 (visited on 10/30/2024).

Leonardi, Nora and Dimitri Van De Ville (Jan. 2015). “On spurious and real fluctuations of dynamic functional connectivity during rest”. In: NeuroImage 104, pp. 430–436. ISSN: 1053-8119. doi: 10.1016/j.neuroimage.2014.09.007. URL: https://www.sciencedirect.com/science/article/pii/S1053811914007496 (visited on 10/30/2024).

Liégeois, Raphaël, Timothy O. Laumann, et al. (Dec. 2017). “Interpreting temporal fluctuations in resting-state functional connectivity MRI”. In: NeuroImage 163, pp. 437–455. ISSN: 1053-8119. doi: 10.1016/j.neuroimage.2017.09.012. URL: https://www.sciencedirect.com/science/article/pii/S1053811917307516 (visited on 11/15/2024).

Liégeois, Raphaël, Jingwei Li, et al. (May 2019). “Resting brain dynamics at different timescales capture distinct aspects of human behavior”. en. In: Nature Communications 10.1. Publisher: Nature Publishing Group, p. 2317. ISSN: 2041-1723. doi: 10.1038/s41467-019-10317-7. URL: https://www.nature.com/articles/s41467-019-10317-7 (visited on 10/26/2024).

Lord, Louis David et al. (2019). “Dynamical exploration of the repertoire of brain networks at rest is modulated by psilocybin”. In: NeuroImage 199.April. Publisher: Elsevier Inc., pp. 127–142. ISSN: 10959572. doi: 10.1016/j.neuroimage.2019.05.060. URL: : https://doi.org/10.1016/j.neuroimage.2019.05.060.

Maeda, E, Hp Robinson, and A Kawana (Oct. 1995). “The mechanisms of generation and propagation of synchronized bursting in developing networks of cortical neurons”. en. In: The Journal of Neuroscience 15.10, pp. 6834–6845. ISSN: 0270-6474, 1529-2401. doi: 10.1523/JNEUROSCI.15-10-06834.1995. URL: https://www.jneurosci.org/lookup/doi/10.1523/JNEUROSCI.15-10-06834.1995 (visited on 11/08/2024).

Mardia, Kanti V. and Peter E. Jupp (May 1999). Directional Statistics. Publication Title: Directional Statistics. John Wiley and Sons Ltd. ISBN: 978-0-470-31697-9. doi: 10.1002/9780470316979. URL: https://onlinelibrary.wiley.com/doi/book/10.1002/9780470316979 (visited on 07/27/2021).

Menon, Vinod (Oct. 2011). “Large-scale brain networks and psychopathology: a unifying triple network model”. In: Trends in Cognitive Sciences 15.10, pp. 483–506. ISSN: 1364-6613. doi: 10.1016/j.tics.2011.08.003. URL: https://www.sciencedirect.com/science/article/pii/S1364661311001719 (visited on 02/04/2025).

ML, Kringelbach et al. (Apr. 2020). “Dynamic coupling of whole-brain neuronal and neurotransmitter systems”. In: Proceedings of the National Academy of Sciences of the United States of America 117.17. Publisher: Proc Natl Acad Sci U S A, pp. 9566–9576. ISSN: 1091-6490. doi: 10.1073/PNAS.1921475117. URL: https://pubmed.ncbi.nlm.nih.gov/32284420/ (visited on 10/22/2021).

Nielsen, Soren F.V. et al. (2017). “Modeling dynamic functional connectivity using a wishart mixture model”. In: 2017 International Workshop on Pattern Recognition in Neuroimaging, PRNI 2017. ISBN: 9781538631591. doi: 10.1109/PRNI.2017.7981505.

Nolte, Guido et al. (Oct. 2004). “Identifying true brain interaction from EEG data using the imaginary part of coherency”. In: Clinical Neurophysiology 115.10, pp. 2292–2307. ISSN: 1388-2457. doi: 10.1016/j.clinph.2004.04.029. URL: https://www.sciencedirect.com/science/article/pii/S1388245704001993 (visited on 11/08/2024).

Olsen, Anders S., Anders Lykkebo-Valløe, et al. (Dec. 2022). “Psilocybin modulation of time-varying functional connectivity is associated with plasma psilocin and subjective effects”. In: NeuroImage 264, p. 119716. ISSN: 1053-8119. doi: 10.1016/j.neuroimage.2022.119716. URL: https://www.sciencedirect.com/science/article/pii/S1053811922008370 (visited on 07/21/2025).

Olsen, Anders S., Jesper D. Nielsen, and Morten Mørup (Aug. 2024). “Coupled Generator Decomposition for Fusion of Electro- and Magnetoencephalography Data”. In: 2024 32nd European Signal Processing Conference (EUSIPCO). ISSN: 2076-1465, pp. 1357–1361. doi: 10.23919/EUSIPCO63174.2024.10715032. URL: https://ieeexplore.ieee.org/document/10715032 (visited on 07/21/2025).

Olsen, Anders S., Emil Ortvald, et al. (June 2023). “Angular Central Gaussian and Watson Mixture Models for Assessing Dynamic Functional Brain Connectivity During a Motor Task”. In: 2023 IEEE International Conference on Acoustics, Speech, and Signal Processing Workshops (ICASSPW), pp. 1–5. doi: 10.1109/ICASSPW59220.2023.10193021. URL: https://ieeexplore.ieee.org/document/10193021 (visited on 04/13/2024).

Pedersen, Mangor et al. (Nov. 2018). “On the relationship between instantaneous phase synchrony and correlation-based sliding windows for time-resolved fMRI connectivity analysis”. In: NeuroImage 181, pp. 85–94. ISSN: 1053-8119. doi: 10.1016/j.neuroimage.2018.06.020. URL: https://www.sciencedirect.com/science/article/pii/S1053811918305275 (visited on 06/03/2024).

Pikovsky, Arkady, Michael Rosenblum, and Jürgen Kurths (2001). Synchronization: A Universal Concept in Nonlinear Sciences. Cambridge Nonlinear Science Series. Cambridge: Cambridge University Press. ISBN: 978-0-521-53352-2. doi: 10.1017/CBO9780511755743. URL: https://www.cambridge.org/core/books/synchronization/E46C1FC3ADC82EEA75AE6F5B9B74E28C (visited on 01/11/2025).

Preti, Maria Giulia, Thomas AW Bolton, and Dimitri Van De Ville (Oct. 2017). “The dynamic functional connectome: State-of-the-art and perspectives”. In: NeuroImage 160. Publisher: Academic Press, pp. 41–54. ISSN: 10959572. doi: 10.1016/j.neuroimage.2016.12.061. (Visited on 07/23/2021).

Rastegarnia, Shima et al. (Dec. 2023). “Brain decoding of the Human Connectome Project tasks in a dense individual fMRI dataset”. In: NeuroImage 283, p. 120395. ISSN: 1053-8119. doi: 10.1016/j.neuroimage.2023.120395. URL: https://www.sciencedirect.com/science/article/pii/S1053811923005463 (visited on 12/08/2025).

Rué-Queralt, Joan et al. (July 2021). “Decoding brain states on the intrinsic manifold of human brain dynamics across wakefulness and sleep”. en. In: Communications Biology 4.1. Publisher: Nature Publishing Group, p. 854. ISSN: 2399-3642. doi: 10.1038/s42003-021-02369-7. URL: https://www.nature.com/articles/s42003-021-02369-7 (visited on 12/01/2025).

Schaefer, Alexander et al. (Sept. 2018). “Local-Global Parcellation of the Human Cerebral Cortex from Intrinsic Functional Connectivity MRI”. In: Cerebral Cortex 28.9. Publisher: Cereb Cortex, pp. 3095–3114. ISSN: 1047-3211. doi: 10.1093/cercor/bhx179. URL: https://pubmed.ncbi.nlm.nih.gov/28981612/ (visited on 09/08/2021).

Shine, James M. et al. (Dec. 2019). “The Low-Dimensional Neural Architecture of Cognitive Complexity Is Related to Activity in Medial Thalamic Nuclei”. In: Neuron 104.5, 849–855.e3. ISSN: 0896-6273. doi: 10.1016/j.neuron.2019.09.002. URL: https://www.sciencedirect.com/science/article/pii/S0896627319307755 (visited on 12/01/2025).

Smith, Stephen M. et al. (Oct. 2013). “Resting-state fMRI in the Human Connectome Project”. In: NeuroImage. Mapping the Connectome 80, pp. 144–168. ISSN: 1053-8119. doi: 10.1016/j.neuroimage.2013.05.039. URL: https://www.sciencedirect.com/science/article/pii/S1053811913005338 (visited on 10/27/2024).

Stevner, A. B. A. et al. (Mar. 2019). “Discovery of key whole-brain transitions and dynamics during human wakefulness and non-REM sleep”. en. In: Nature Communications 10.1. Publisher: Nature Publishing Group, p. 1035. ISSN: 2041-1723. doi: 10.1038/s41467-019-08934-3. URL: https://www.nature.com/articles/s41467-019-08934-3 (visited on 10/23/2024).

Sun, Junfeng and Michael Small (Oct. 2009). “Unified framework for detecting phase synchronization in coupled time series”. In: Physical Review E 80.4. Publisher: American Physical Society, p. 046219. doi: 10.1103/PhysRevE.80.046219. URL: https://link.aps.org/doi/10.1103/PhysRevE.80.046219 (visited on 01/11/2025).

Thomas Yeo, B. T. et al. (Sept. 2011). “The organization of the human cerebral cortex estimated by intrinsic functional connectivity”. In: Journal of Neurophysiology 106.3. Publisher: American Physiological Society, pp. 1125–1165. ISSN: 0022-3077. doi: 10.1152/jn.00338.2011. URL: https://journals.physiology.org/doi/full/10.1152/jn.00338.2011 (visited on 11/15/2024).

Tian, Ye et al. (Nov. 2020). “Topographic organization of the human subcortex unveiled with functional connectivity gradients”. en. In: Nature Neuroscience 23.11. Publisher: Nature Publishing Group, pp. 1421–1432. ISSN: 1546-1726. doi: 10.1038/s41593-020-00711-6. URL: https://www.nature.com/articles/s41593-020-00711-6 (visited on 05/15/2024).

Tyler, David E. (Sept. 1987). “Statistical Analysis for the Angular Central Gaussian Distribution on the Sphere”. In: Biometrika 74.3. Publisher: JSTOR, p. 579. ISSN: 00063444. doi: 10.2307/2336697. (Visited on 10/21/2022).

Uhlhaas, Peter et al. (July 2009). “Neural synchrony in cortical networks: history, concept and current status”. English. In: Frontiers in Integrative Neuroscience 3. Publisher: Frontiers. ISSN: 1662-5145. doi: 10.3389/neuro.07.017.2009. URL: https://www.frontiersin.org/journals/integrative-neuroscience/articles/10.3389/neuro.07.017.2009/full (visited on 11/15/2024).

Van Essen, D. C. et al. (Oct. 2012). “The Human Connectome Project: a data acquisition perspective”. eng. In: NeuroImage 62.4, pp. 2222–2231. ISSN: 1095-9572. doi: 10.1016/j.neuroimage.2012.02.018.

Van Essen, David C. et al. (Oct. 2013). “The WU-Minn Human Connectome Project: an overview”. In: NeuroImage 80. Publisher: Neuroimage, pp. 62–79. ISSN: 1095-9572. doi: 10.1016/J.NEUROIMAGE.2013.05.041. URL: https://pubmed.ncbi.nlm.nih.gov/23684880/ (visited on 10/20/2022).

Vidaurre, Diego, Andrew J. Quinn, et al. (Feb. 2016). “Spectrally resolved fast transient brain states in electrophysiological data”. In: NeuroImage 126, pp. 81–95. ISSN: 1053-8119. doi: 10.1016/j.neuroimage.2015.11.047. URL: https://www.sciencedirect.com/science/article/pii/S1053811915010691 (visited on 10/26/2024).

Vidaurre, Diego, Stephen M. Smith, and Mark W. Woolrich (2017). “Brain network dynamics are hierarchically organized in time”. In: Proceedings of the National Academy of Sciences of the United States of America 114.48, pp. 12827–12832. ISSN: 10916490. doi: 10.1073/pnas.1705120114.

Vohryzek, Jakub et al. (Apr. 2020). “Ghost Attractors in Spontaneous Brain Activity: Recurrent Excursions Into Functionally-Relevant BOLD Phase-Locking States”. In: Frontiers in Systems Neuroscience 14. Publisher: Frontiers, p. 20. ISSN: 16625137. doi: 10.3389/fnsys.2020.00020. (Visited on 10/22/2021).

Wang, Huifang E. et al. (Dec. 2014). “A systematic framework for functional connectivity measures”. English. In: Frontiers in Neuroscience 8. Publisher: Frontiers. ISSN: 1662-453X. doi: 10.3389/fnins.2014.00405. URL: https://www.frontiersin.org/journals/neuroscience/articles/10.3389/fnins.2014.00405/full (visited on 10/23/2024).

Wang, Xiaoxiao et al. (Dec. 2019). “Decoding and mapping task states of the human brain via deep learning”. In: Human Brain Mapping 41.6, pp. 1505–1519. ISSN: 1065-9471. doi: 10.1002/hbm.24891. URL: https://pmc.ncbi.nlm.nih.gov/articles/PMC7267978/ (visited on 12/08/2025).

Womelsdorf, Thilo et al. (June 2007). “Modulation of Neuronal Interactions Through Neuronal Synchronization”. In: Science 316.5831. Publisher: American Association for the Advancement of Science, pp. 1609–1612. doi: 10.1126/science.1139597. URL: https://www.science.org/doi/10.1126/science.1139597 (visited on 11/15/2024).

Zarghami, Tahereh S., Gholam-Ali Hossein-Zadeh, and Fariba Bahrami (Mar. 2020). “Deep Temporal Organization of fMRI Phase Synchrony Modes Promotes Large-Scale Disconnection in Schizophrenia”. English. In: Frontiers in Neuroscience 14. Publisher: Frontiers. ISSN: 1662-453X. doi: 10.3389/fnins.2020.00214. URL: https://www.frontiersin.org/journals/neuroscience/articles/10.3389/fnins.2020.00214/full (visited on 10/13/2024).

Zhang, Yu et al. (May 2021). “Functional annotation of human cognitive states using deep graph convolution”. In: NeuroImage 231, p. 117847. ISSN: 1053-8119. doi: 10.1016/j.neuroimage.2021.117847. URL: https://www.sciencedirect.com/science/article/pii/S1053811921001245 (visited on 12/14/2025).

