## Supporting information for "Uncovering dynamic human brain phase coherence networks"

#### 1 Notes on the complex angular central Gaussian distribution

The complex angular central Gaussian (CACG) distribution density is given by ([1])

$$f_{\mathrm{CACG}}(\mathbf{u};\boldsymbol{\Psi}) = \frac{\Gamma(p)}{2\pi^p|\boldsymbol{\Psi}|}\left(\mathbf{u}^{\mathrm{H}}\boldsymbol{\Psi}^{-1}\mathbf{u}\right)^{-p},\ \mathbf{u}\in\mathbb{CP}^{p-1}.$$

Here, $\boldsymbol{\Psi}\in\mathbb{C}^{p\times p}$ is a symmetric positive-definite matrix identifiable up to a positive scale factor, and $|\cdot|$ is the matrix determinant. $\boldsymbol{\Psi}$ is typically normalized to have $\mathrm{tr}(\boldsymbol{\Psi}) = p$ for convenience due to the scale-invariance of the density.

##### 1.1 Maximum likelihood estimation

The log-likelihood of the ACG model is

$$l(\boldsymbol{\Psi};\mathbf{u}) = -\log|\boldsymbol{\Psi}| - p\log\left(\mathbf{u}^{\mathrm{H}}\boldsymbol{\Psi}^{-1}\mathbf{u}\right) + \delta,$$

where $\delta$ indicates terms unrelated to $\boldsymbol{\Psi}$. The maximum likelihood solution for $n > p(p-1)$ is ([2])

$$\boldsymbol{\Psi} = p\frac{\mathbf{u}\mathbf{u}^{\mathrm{H}}}{\mathbf{u}^{\mathrm{H}}\boldsymbol{\Psi}^{-1}\mathbf{u}}.$$

The above ML estimator can be approached using an iterative fixed point algorithm. Such an algorithm must necessarily evaluate convergence of $\boldsymbol{\Psi}$, which may be measured using the Frobenius norm of the variable in consecutive iterations $j$ and $j-1$:

$$\mathcal{L} = ||\boldsymbol{\Psi}_j - \boldsymbol{\Psi}_{j-1}||_F^2$$

Since $\boldsymbol{\Psi}$ is scale-invariant, the Frobenius norm only makes sense if the estimate at consecutive iterations are of the same scale. Thus, to track convergence of the fixed point update, trace-normalization is necessary. Since $\mathrm{tr}(\mathbf{u}_t\mathbf{u}_t^{\mathrm{H}}) = 1$ the trace may be simplified to

$$\mathrm{tr}(\boldsymbol{\Psi}) = \frac{p}{n}\sum_{t=1}^{n}\frac{1}{\mathbf{u}_t^{\top}\boldsymbol{\Psi}^{-1}\mathbf{u}_t}.$$

Thus, the estimate at each iteration of the fixed point algorithm is scaled by the trace before evaluating convergence.

#### 1.2 Low-rank-plus-diagonal approximation

$\boldsymbol{\Psi}$ may be approximated using a low-rank-plus-diagonal reparameterization: $\boldsymbol{\Psi} \approx \mathbf{M}\mathbf{M}^{\mathrm{H}} + \mathbf{I}_p = \mathbf{Z}$, where $\mathbf{M} \in \mathbb{C}^{p\times r}$. Here, $r \leq p$ is the rank modeled and the identity matrix ensures stability of the determinant and invertibility of $\boldsymbol{\Psi}$. Using standard matrix calculus (see, e.g., the Matrix Cookbook), we find the maximum likelihood estimator to be

$$\mathbf{M} = p\frac{\mathbf{u}\mathbf{u}^{\top}\mathbf{Z}^{-1}\mathbf{M}}{\mathbf{u}^{\top}\mathbf{Z}^{-1}\mathbf{u}}.$$

Notably, Woodbury's matrix inversion lemma can be utilized for faster inversion of $\mathbf{Z}$. We introduce $\mathbf{D} = \mathbf{M}^{\top}\mathbf{M} + \mathbf{I}_r$, such that

$$\begin{aligned}\mathbf{Z}^{-1} &= \left(\mathbf{M}\mathbf{M}^{\mathrm{H}} + \mathbf{I}_p\right)^{-1} \\ &= \mathbf{I}_p - \mathbf{M}\mathbf{D}^{-1}\mathbf{M}^{\mathrm{H}}.\end{aligned}$$

The log-likelihood function is then given by:

$$l(\mathbf{M};\mathbf{u}) = -\log|\mathbf{Z}| - p\log\left(1 - \mathbf{u}^{\mathrm{H}}\mathbf{M}\mathbf{D}^{-1}\mathbf{M}^{\mathrm{H}}\mathbf{u}\right) + \delta$$

and the maximum likelihood estimator:

$$\mathbf{M} = p\frac{\mathbf{u}\mathbf{u}^{\mathrm{H}}\mathbf{M}\mathbf{D}^{-1}}{1 - \mathbf{u}^{\mathrm{H}}\mathbf{M}\mathbf{D}^{-1}\mathbf{M}^{\mathrm{H}}\mathbf{u}}.$$

To ensure that the lowrank-plus-diagonal approximation is also trace-normalized we introduce a scalar factor $\tilde{\mathbf{Z}} = \gamma\mathbf{Z}$, where

$$\begin{aligned}\gamma &= \frac{p}{\mathrm{tr}\,(\mathbf{Z})} \\ &= \frac{1}{o/p + 1}\end{aligned}$$

with $o = ||\mathbf{M}||_F^2$. The maximum-likelihood estimate for $\mathbf{M}$ given does not change upon introducing $\gamma$. Thus, we introduce a rescaling $\tilde{\mathbf{M}} = \sqrt{\gamma}\mathbf{M}$ for the $\mathbf{M}$-update while only including the scaling by $\gamma$ on the identity matrix, i.e., $\tilde{\mathbf{Z}} = \tilde{\mathbf{M}}\tilde{\mathbf{M}}^{\mathrm{H}} + \gamma\mathbf{I}$.

### 2 Hidden Markov Models

Mixture models do not necessarily capture the sequential dependencies present in ordered data, such as a multivariate time-series. Notably, Hidden Markov Models (HMMs) explicitly model transition dynamics between components by imposing a transition probability matrix $\mathbf{T}$ with elements $T_{k',k}$, which governs the transition probability from state $k'$ to state $k$. Given the complexity of EM optimization for HMMs, especially for large data sets, we instead utilized the direct numerical optimization method of the log-likelihood function using the forward algorithm:

$$\alpha_{t,k}^{(s)} = \sum_{k'=1}^{K} \alpha_{t-1,k'}^{(s)} T_{k',k} f(\mathbf{y}_t^{(s)}|\boldsymbol{\Psi}_k),$$

where $\alpha_{t,k}^{(s)}$ represents the probability of being in state $k$ at time $t$ for the $s^{\mathrm{th}}$ sequence (e.g., a subject or separate recording), and $\alpha_{1,k}^{(s)} = w_k f(\mathbf{y}_1^{(s)}|\boldsymbol{\Psi}_k)$. Of note, the -parameter for HMM only applies to the first sample in each sequence and thus has a different interpretation than for mixture models. The log-likelihood to be maximized numerically is

$$\mathcal{L}_{\text{HMM}}(\boldsymbol{\Psi}, \mathbf{w}, \mathbf{T}|\mathcal{Y}) = \sum_{s=1}^{S} \ln \left( \sum_{k=1}^{K} \alpha_{n_s,k}^{(s)} \right),$$

where $S$ is the number of observation sequences, and $n_s$ is the length of sequence $s$. Following the parameter optimization, we used the Viterbi decoder [3] to extract state assignments.

### 3 Normalized Mutual Information

Given two binary or weighted responsibility matrices $\boldsymbol{\beta}_1$ and $\boldsymbol{\beta}_2$ of size $K_1 \times N$ and $K_2 \times N$, respectively, which sum to one for each column, normalized mutual information (NMI) is defined as [4]

$$\begin{aligned}
\text{NMI}(\boldsymbol{\beta}_1, \boldsymbol{\beta}_2) &= \frac{2\text{MI}(\boldsymbol{\beta}_1, \boldsymbol{\beta}_2)}{\text{MI}(\boldsymbol{\beta}_1, \boldsymbol{\beta}_1) + \text{MI}(\boldsymbol{\beta}_2, \boldsymbol{\beta}_2)}, \\
\text{MI}(\boldsymbol{\beta}_1, \boldsymbol{\beta}_2) &= \sum_{k_1,k_2} p(k_1, k_2) \log \frac{p(k_1, k_2)}{p(k_1)p(k_2)}, \\
p(k_1, k_2) &= \frac{1}{N} \sum_{n} \beta_{k_1,n} \beta_{k_2,n}.
\end{aligned}$$

### 4 Supporting figures and tables

In this section we include additional information and supporting experimentation.

- Table S1 outlines additional properties of the considered statistical distributions.
- Figure S1 describes the synthetic generation of three-dimensional data points on the projective hyperplane and results from clustering these with varying models.
- Figure S2 shows modeling results on phase-controlled resting-state fMRI.
- Figure S3 extends Figure S3 by also showing varying initialization method and estimation method (EM, stochastic, HMM).
- Figure S4 presents clustering performance using all models on task fMRI data when varying parcellation size, excluding global signal regression (GSR), and HCP dataset version.
- Figure S5 presents clustering performance using all models on task fMRI data in 116 parcels following GSR when varying training data split.
- Figure S6 shows the main results (Fig. 4) where all task volumes, including pre-stimulus volumes, were used for training, and models were evaluated on all samples, including inter-trial intervals.
- Figure S7 shows the average test-set posterior probability (average over 99 subjects) for the Gaussian model (for amplitude coupling) trained on task-fMRI data.
- Figure S8 shows the average test-set posterior probability (average over 99 subjects) for the complex Gaussian model (for phase-amplitude coupling) trained on task-fMRI data.
- Figure S9 shows the resting state results for the Gaussian model
- Figure S10 shows the resting state results for the complex Gaussian model
- Figure S11 presents the global phase coherence matrix before and after global signal regression, as well as single-component estimates for all statistical distributions.
- Figure S12 shows K-means performance compared to the initialization method.

Table S1: Overview of clustering methods. Models in blue indicate methods not previously applied to functional neuroimaging data. PH: Projective hyperplane.

| | **Complex** | **Anisotropic** | **Probabilistic** | **Manifold** |
|---|---|---|---|---|
| *Time-series* | | | | |
| **Gaussian mixture** | ✗ | ✓ | ✓ | Euclidean |
| *Analytic signal* | | | | |
| **Complex Gaussian mixture** | ✓ | ✓ | ✓ | Complex Euclidean |
| *Complex dynamic phase coherence maps* | | | | |
| **Complex angular central Gaussian mixture** | ✓ | ✓ | ✓ | Complex PH |
| **Complex diametrical clustering** | ✓ | ✗ | ✗ | Complex PH |
| *Leading eigenvector of cosine phase coherence maps (LEiDA)* | | | | |
| **Diametrical clustering** | ✗ | ✗ | ✗ | PH |
| **Least squares K-means (sign-flipped eigenvectors)** | ✗ | ✗ | ✗ | Euclidean |

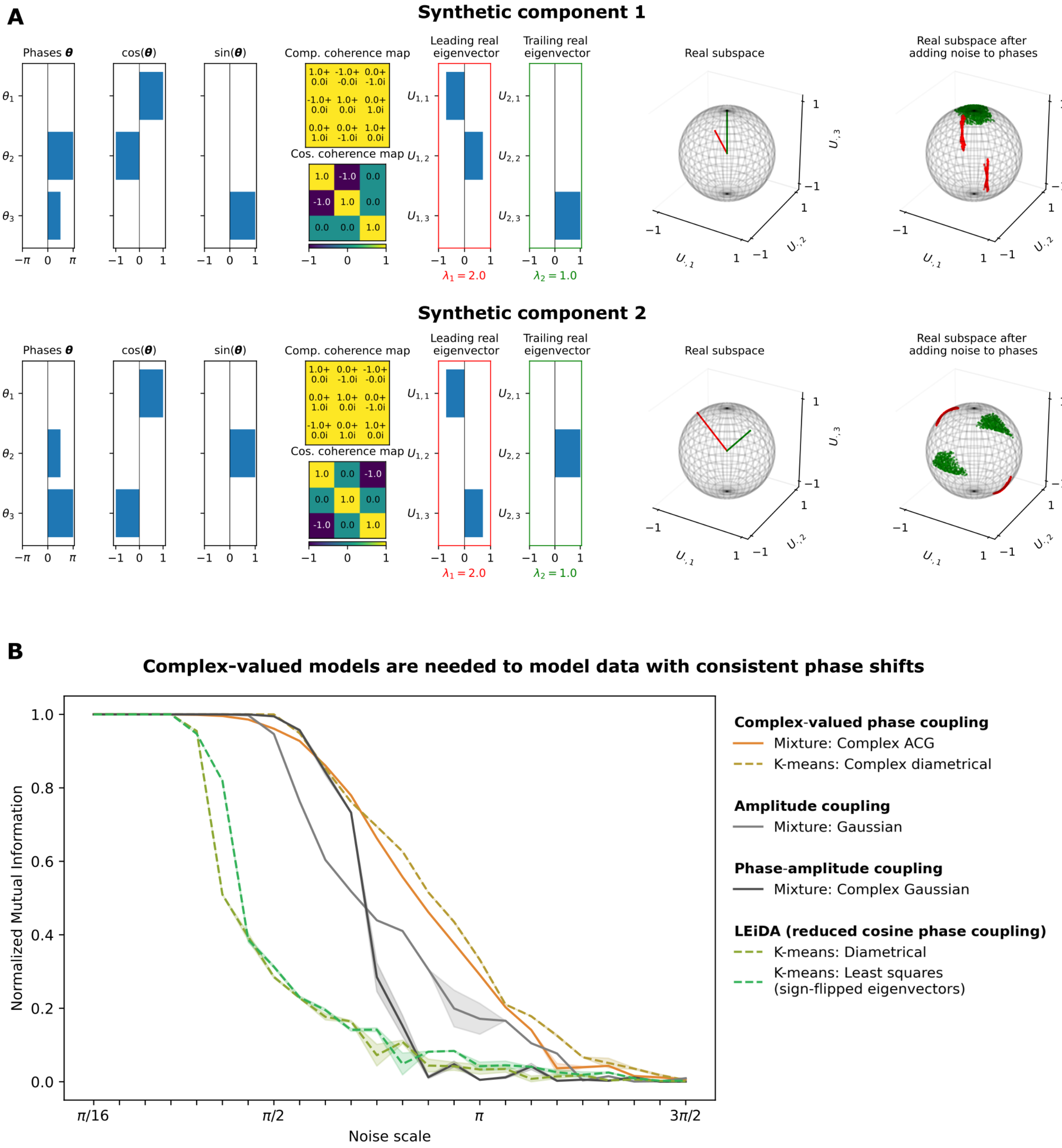


Figure S1: Model performance on synthetic data with consistent phase shifts. A) Synthetic generation of 3-dimensional data points. An initial phase vector is defined, with corresponding complex or cosine phase coherence matrix and eigendecomposition. Train and test sets were generated identically with $n = 1000$ points for each component. B) Clustering performance measured by normalized mutual information between ground-truth and predicted labels (K-means, dashed lines) or posterior probability (probabilistic mixtures, solid lines) on the test set. Shaded areas represent the standard error across 5 initializations; probabilistic mixture models were initialized using their corresponding K-means method, which was seeded using K-means++. All covariance matrices were reparameterized with a rank-1 matrix (complex ACG) or a rank-2 matrix (Gaussian, complex Gaussian) to correspond with the data's rank.

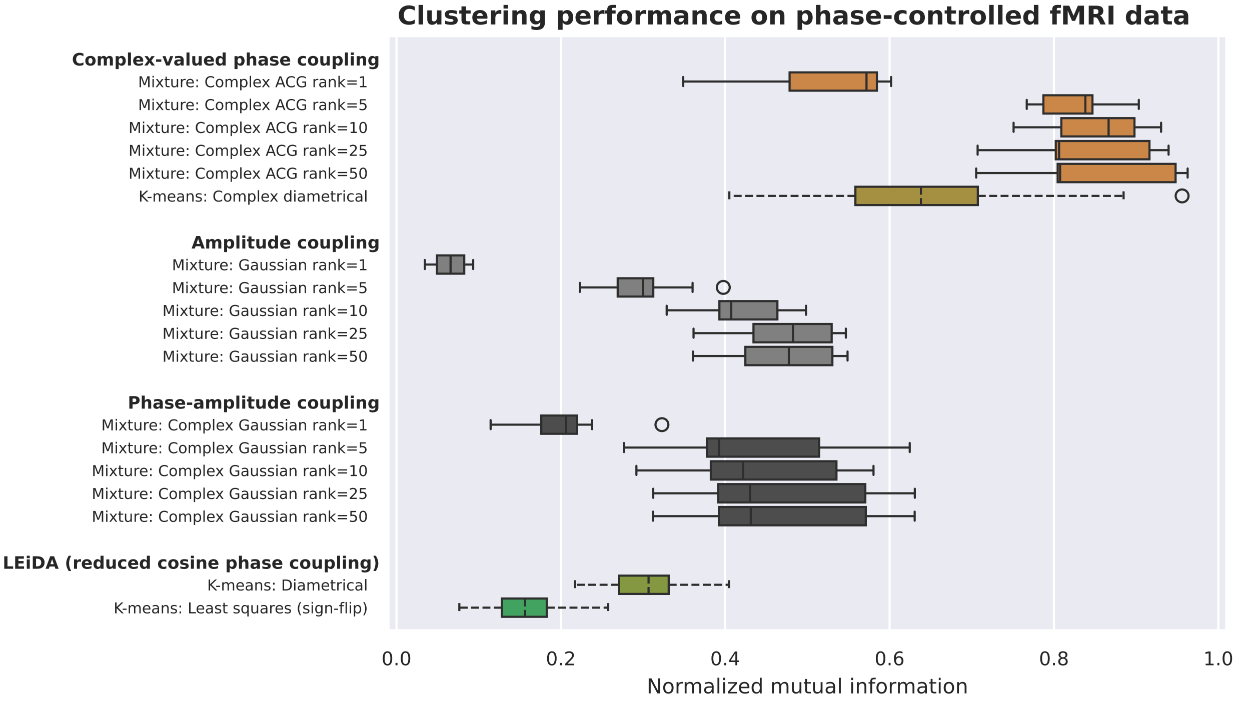


Figure S2: Clustering of phase-controlled 116-region fMRI data from a single subject. Model rank was increased gradually and rank-1 models initialized using K-means clustering seeded using K-means++. Optimization was performed using PyTorch (green boxes in Supplementary Fig. S3). Normalized mutual information between true labels and the estimated train data partition or posterior probability from 10 initializations are shown.

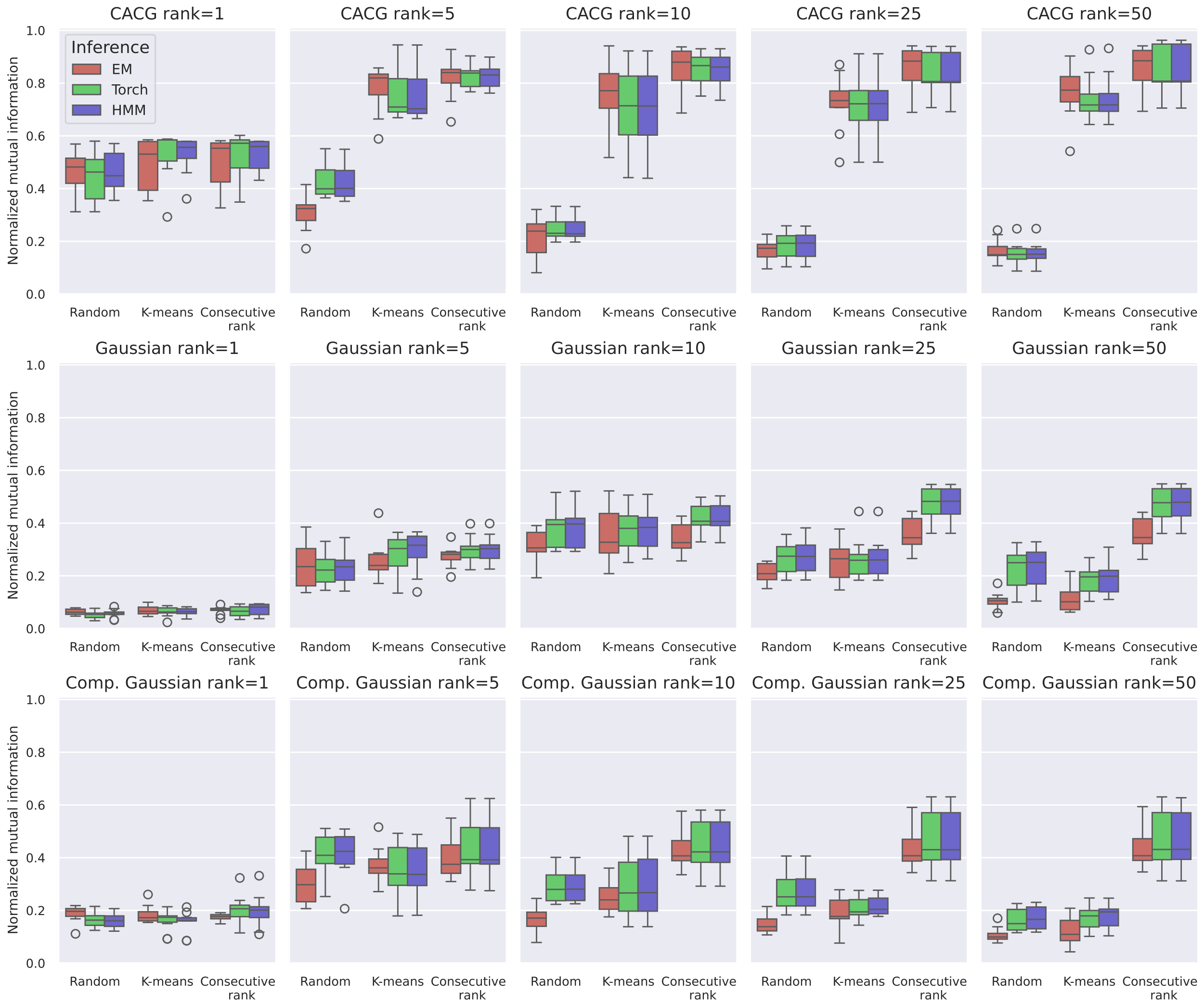


Figure S3: Effect of initialization and model estimation method. "Consecutive rank" represents the scenario, where higher-rank models are initialized from lower rank models. Hidden Markov models (HMM) were always initialized from the PyTorch mixture estimate of the same rank.

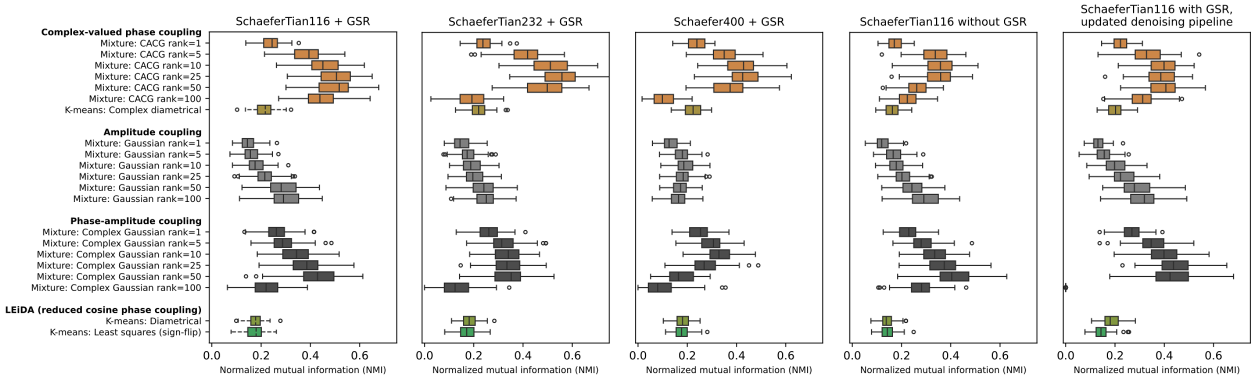


Figure S4: Mixture modeling performance depending on parcellations, global signal regression (GSR), and human connectome project denoising pipeline version. The 116-dimensional Schaefer-Tian atlas ([5, 6]) with GSR (left-most graph) are presented in the main paper. This corresponded to the 2018-version of the HCP dataset, while the right-most graph corresponded to a 2025-version including temporal ICA artifact removal. Normalized mutual information (NMI) between true scan type and test set posterior mixture model probability across 99 unseen subjects is shown. Each model was run 10 times and only the best model, in terms of train-set NMI, is shown.

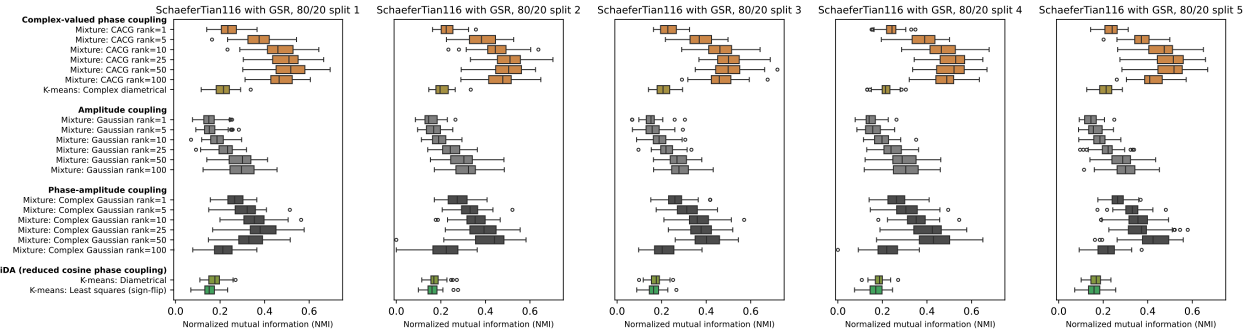


Figure S5: Mixture modeling performance depending on dataset partition. In the main analysis, 155 subjects were used for training while 99 subjects were used for testing. In order to test whether the specific dataset partition influenced results, we performed an additional analysis splitting the 254 subjects into 5 groups (51 subjects in each group). In each split, one group was held out for testing while the remaining four groups were used for training the models. Normalized mutual information (NMI) between true scan type and test set posterior mixture model probability across the 51 unseen subjects is shown. Each model was run 5 times and only the best model, in terms of train-set NMI, is shown.

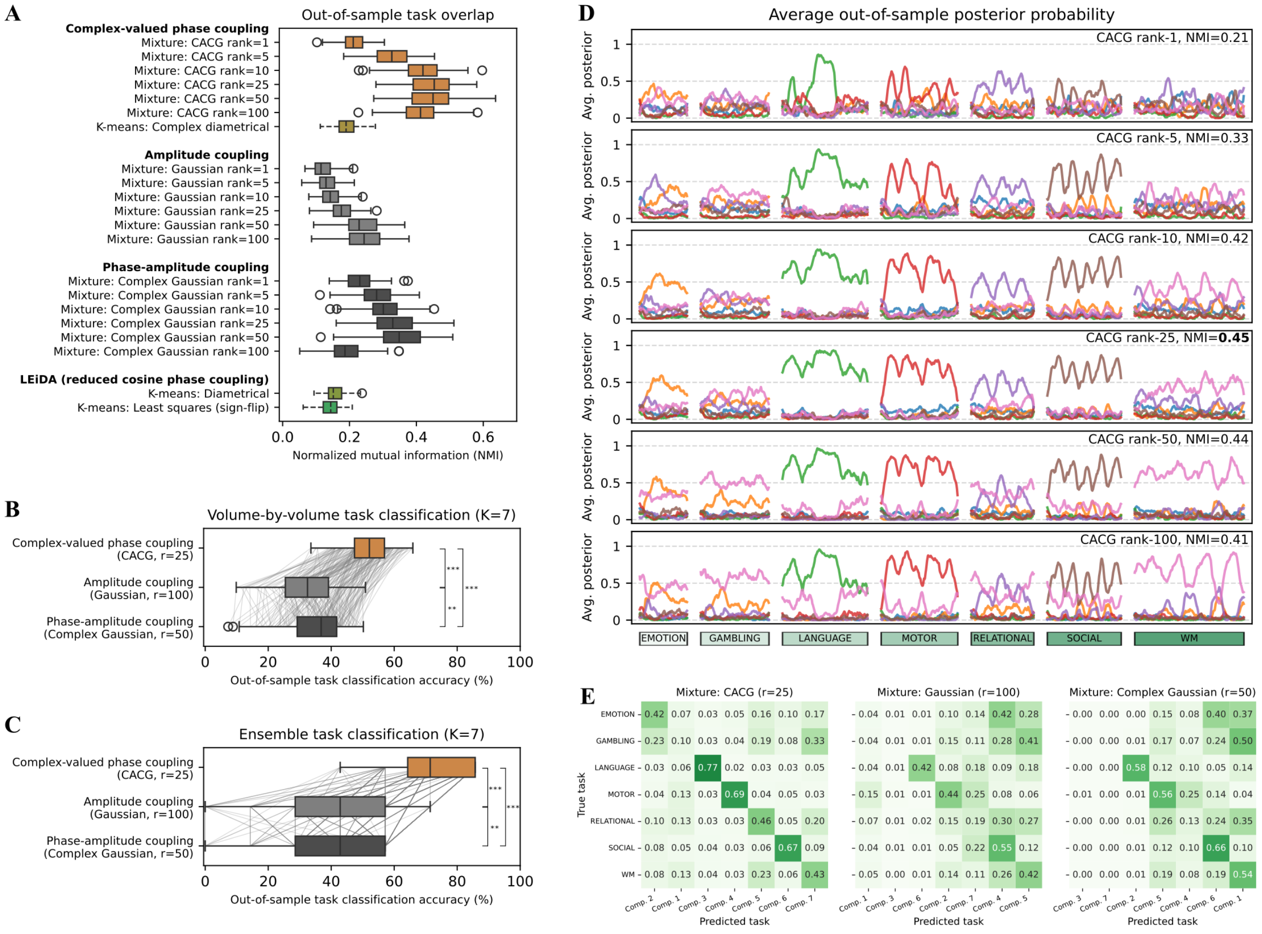


Figure S6: Seven-component mixture models applied to the seven task fMRI scans from the human connectome project (HCP), including pre-stimulus volumes and evaluating accuracy in resting blocks as well. **A**): Overlap between posterior probability and the true task labels measured using normalized mutual information. **B**): Volume-by-volume task classification accuracy using binarized test set posterior probabilities. Components were matched to tasks based on their average performance across train subjects. The average classification accuracy across volumes and scans is shown. **C**): Same as B), with only one predicted component per scan based on the average within-scan posterior. In A), B), and C), boxplots depict values for N=99 test subjects. In B) and C), ** $p < 0.01$, *** $p < 0.001$ indicates significance in a Wilcoxon signed rank test, Bonferroni-corrected for three comparisons. **D**): The average test posterior of each of the seven components (one color for each) of the CACG mixture model at different component ranks. The x-axis represents time with whitespace demarcating different task scans. Boxes indicate the name of the task and box widths are proportional to the number of volumes in the task scan. **E**): Test-set confusion matrices after ordering components to their most likely task as measured by train accuracy.

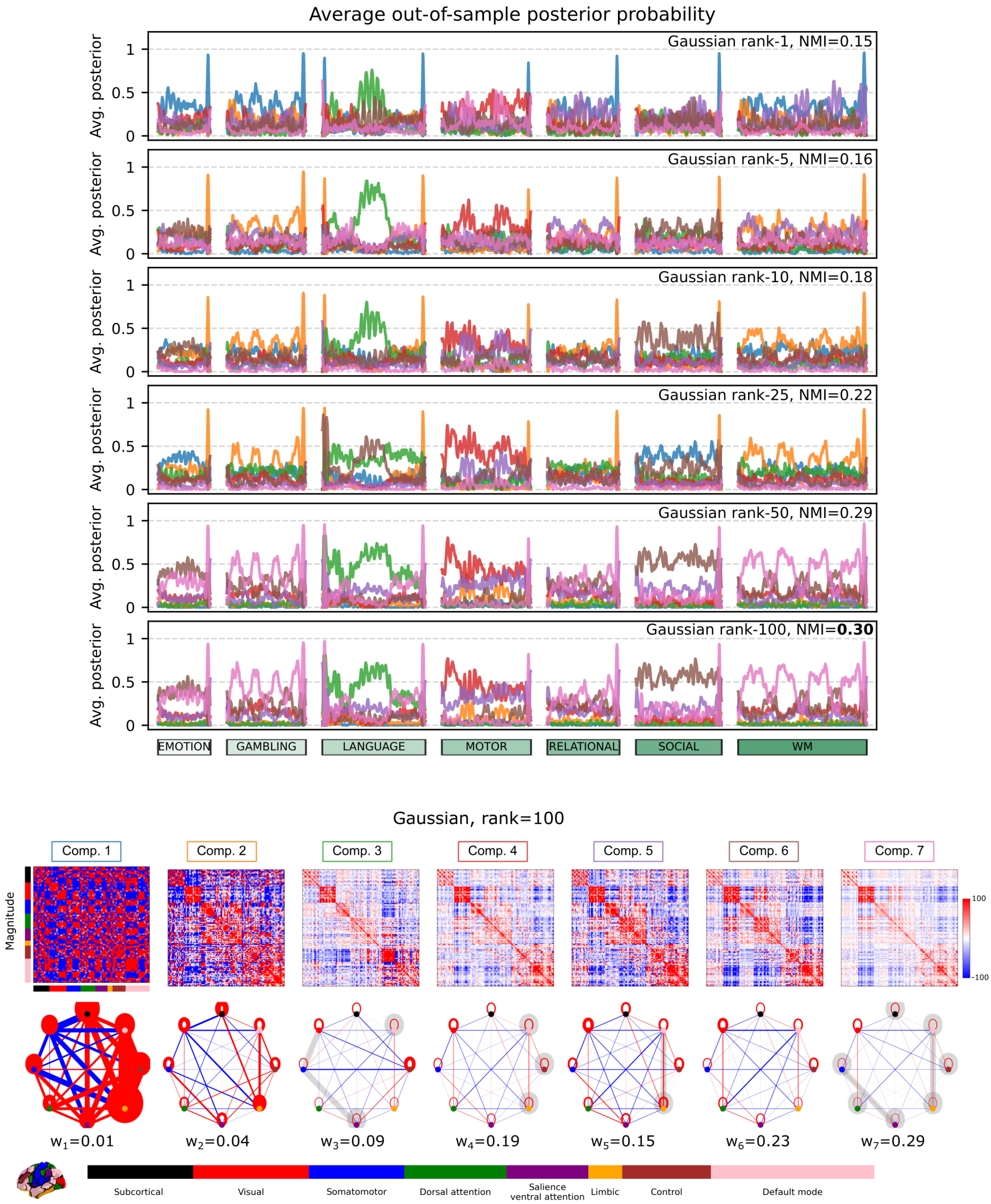


Figure S7: The average test posterior and components of the Gaussian mixture ($K = 7$) at different ranks. The model was trained on fMRI data from 7 tasks from 155 subjects from the Human connectome project data. Normalized mutual information (NMI) indicates the overlap between the test posterior probability and the true task labels computed for each of 99 test subjects and averaged across subjects.

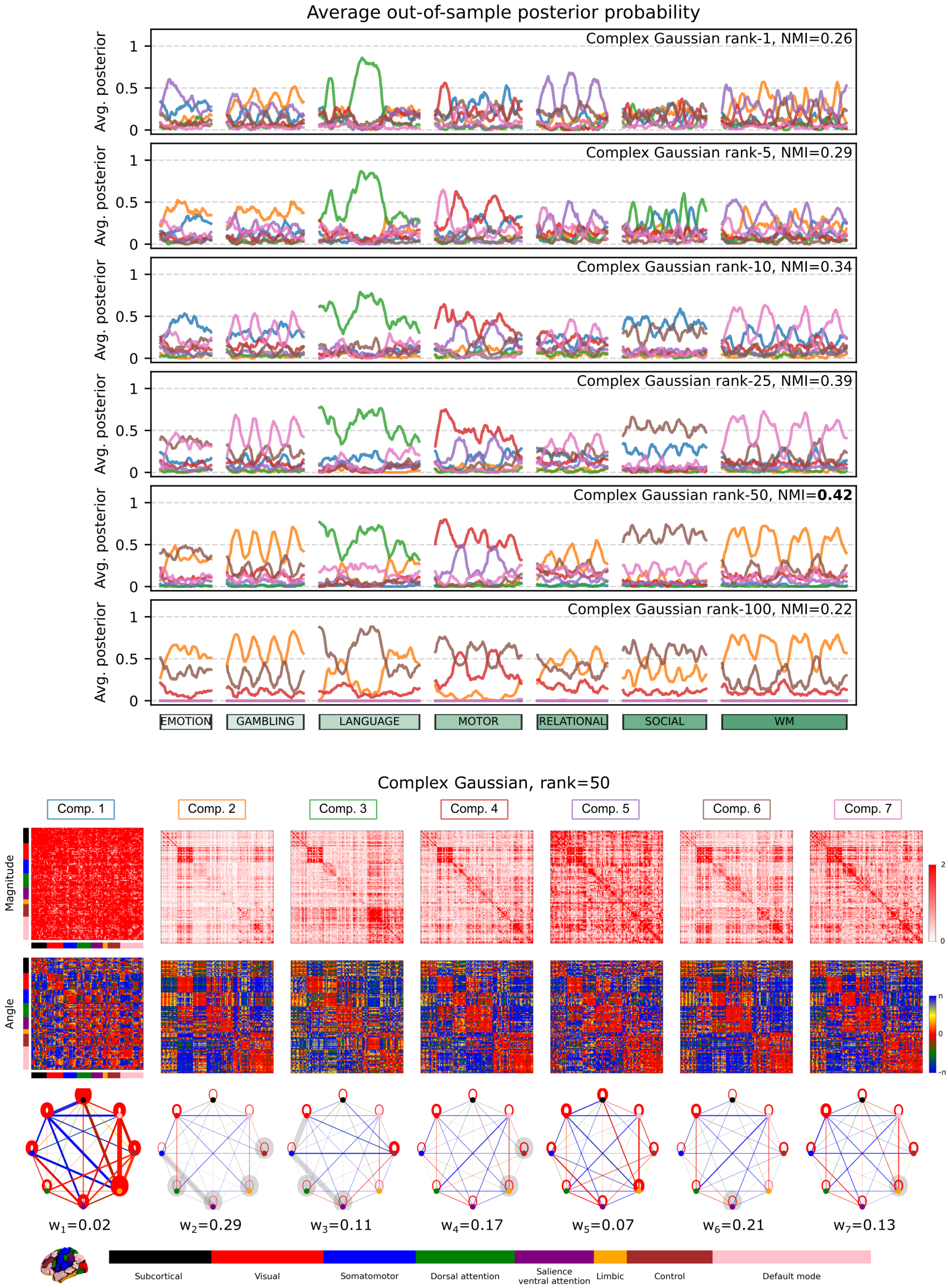


Figure S8: The average test posterior and components of the complex Gaussian mixture ($K = 7$) at different ranks. The model was trained on fMRI data from 7 tasks from 155 subjects from the Human connectome project data. Normalized mutual information (NMI) indicates the overlap between the test posterior probability and the true task labels computed for each of 99 test subjects and averaged across subjects.

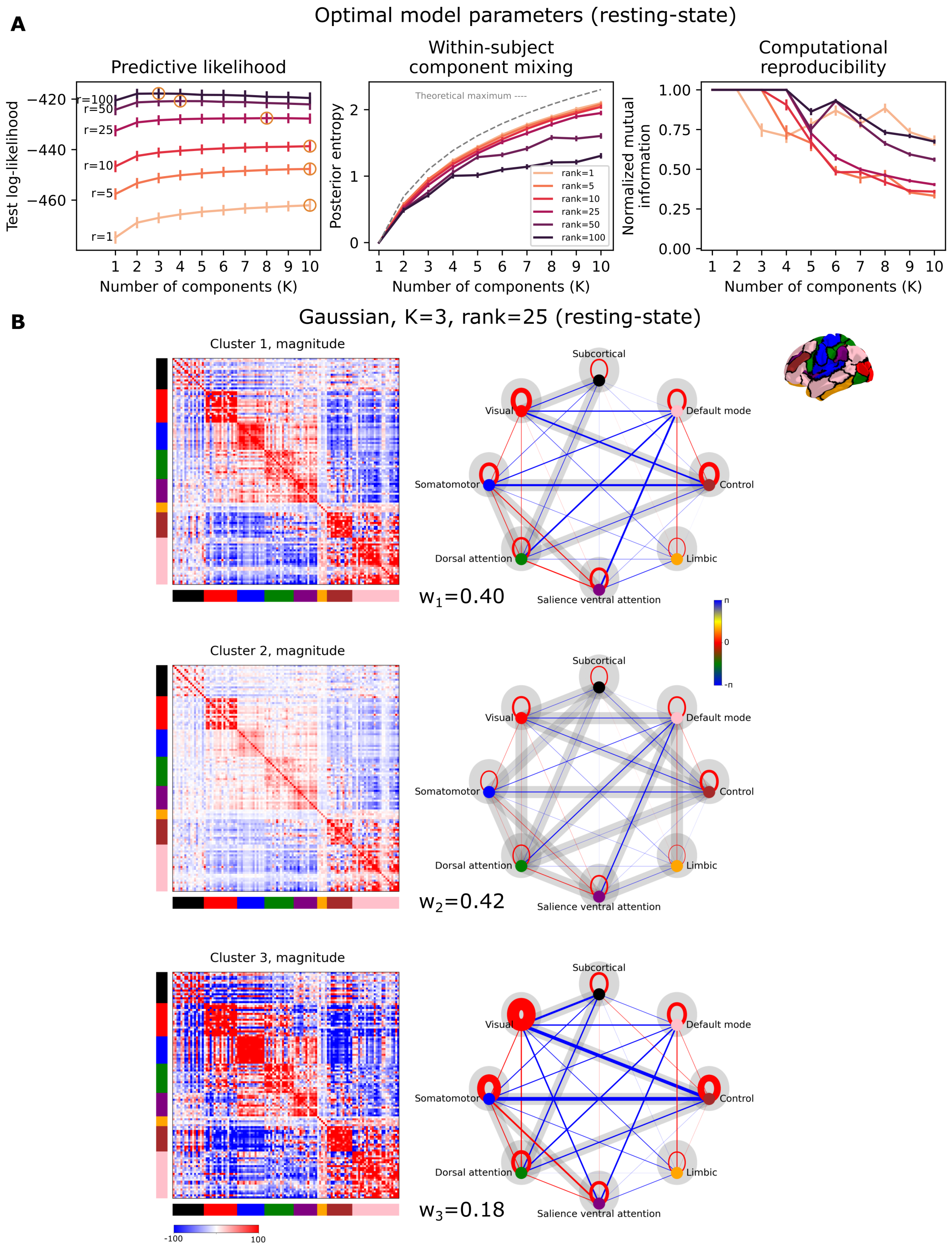


Figure S9: Resting-state results for the Gaussian mixture model.

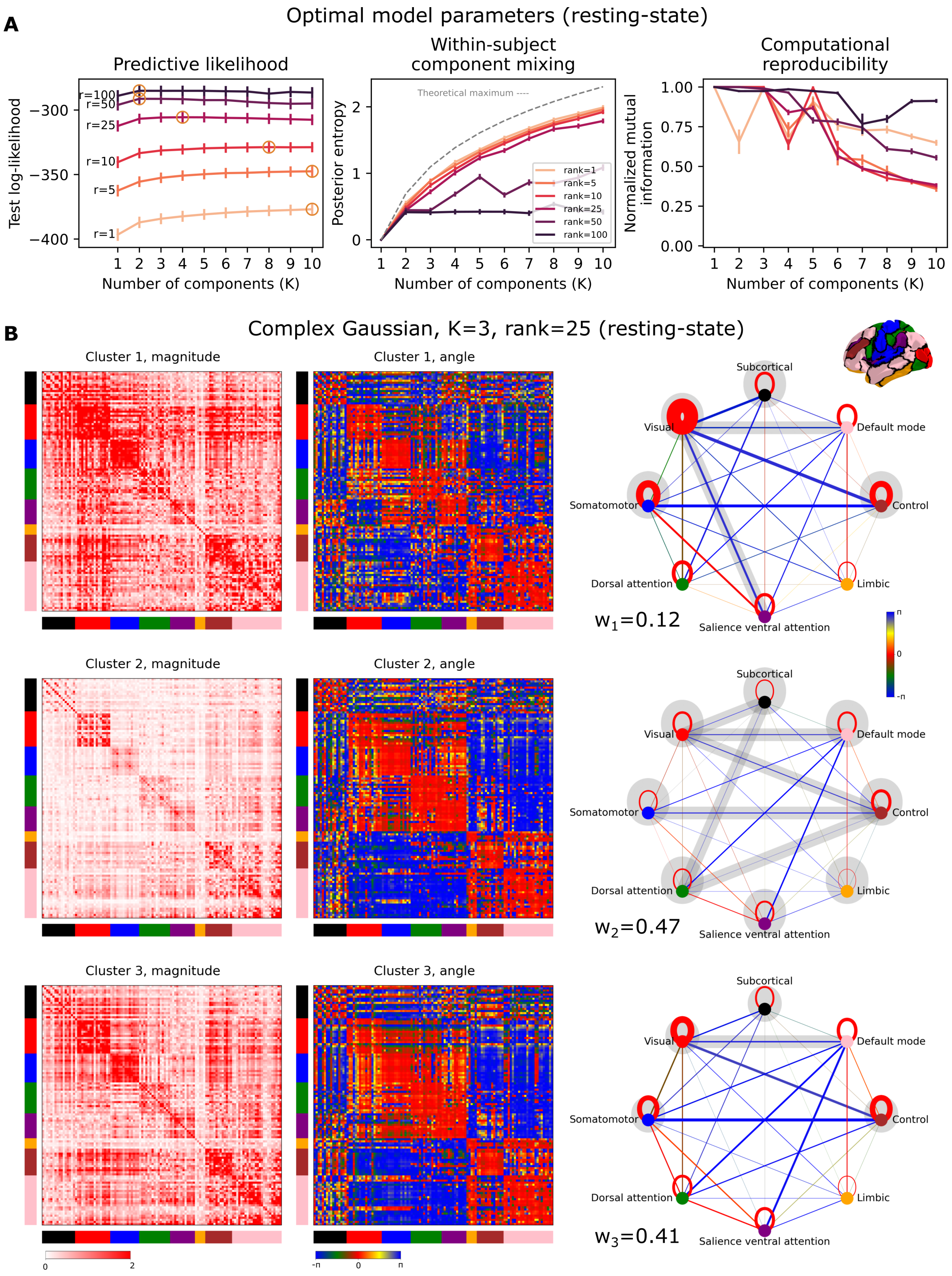


Figure S10: Resting-state results for the Complex Gaussian mixture model.

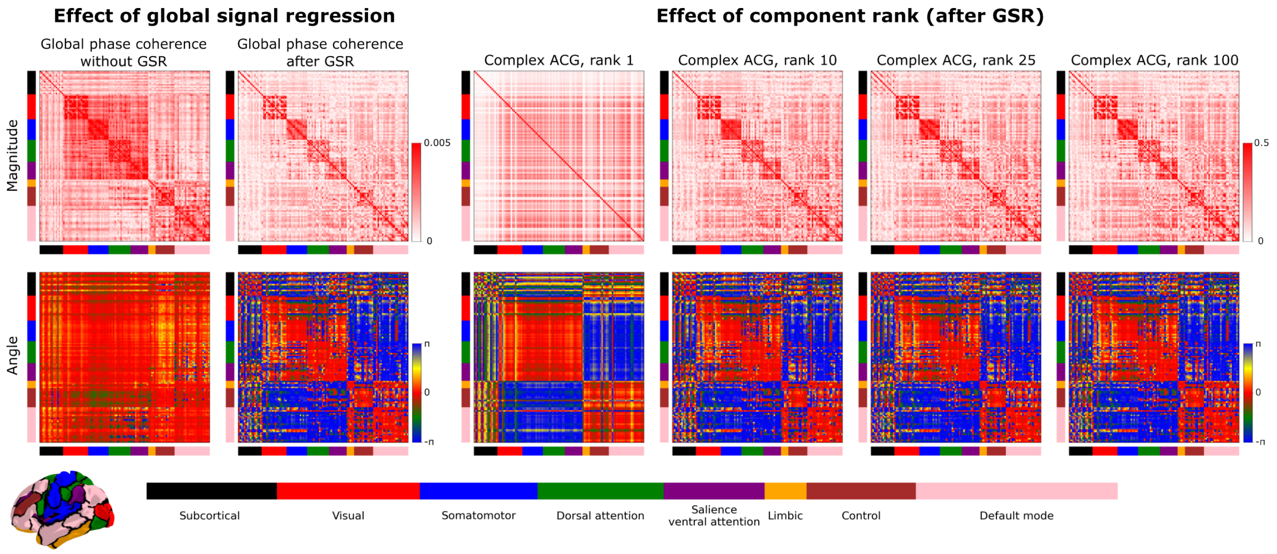


Figure S11: Estimation of global resting-state phase coherence maps from 255 subjects in the 116-dimensional Schaefer-Tian atlas [5, 6]. A): The global average complex phase coherence matrix with and without global signal regression (GSR); the magnitude is also known as the phase-locking value. B): Single-component CACG shown as $\mathbf{Z} = \mathbf{M}\mathbf{M}^{\mathrm{H}} + \mathbf{I}$ estimates on data, where the global signal had been regressed out. The colored boxes indicate a partition of the 116 areas into 8 networks defined by the 7 Yeo networks and an added subcortical network.

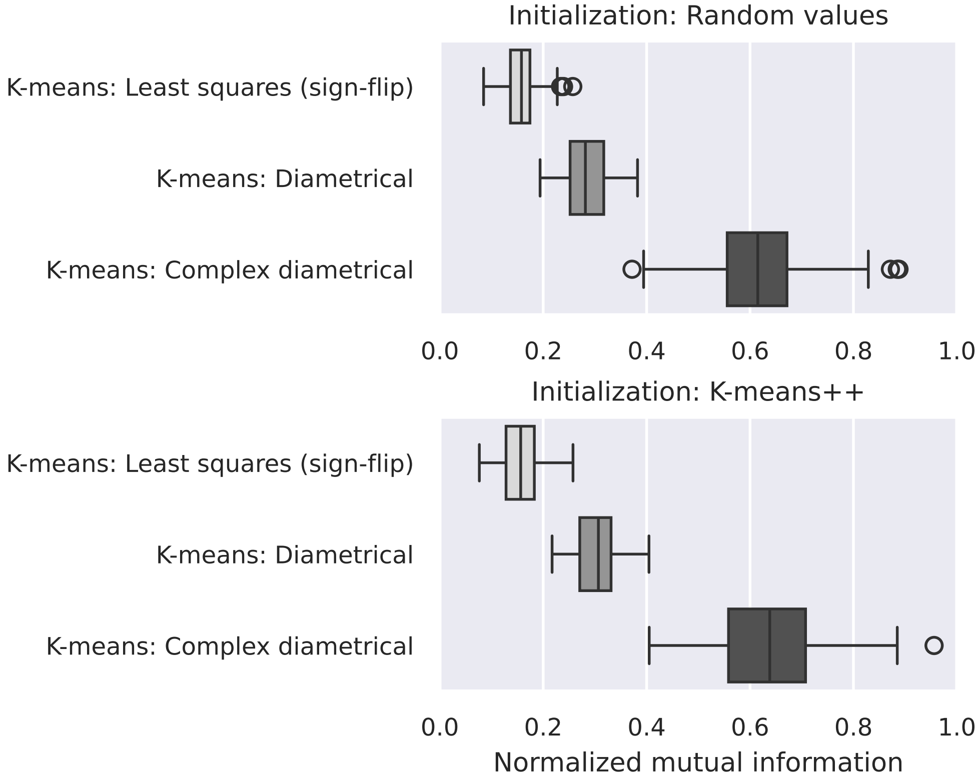


Figure S12: Effect of initialization strategy on Riemannian K-means clustering over 25 repeats of each clustering algorithm with either random uniform centroids or centroids chosen from the dataset using ++-seeding.

### References


[1] John T. Kent. “Data analysis for shapes and images”. In: *Journal of Statistical Planning and Inference*. Robust Statistics and Data Analysis, Part II 57.2 (Feb. 1997), pp. 181–193. ISSN: 0378-3758. DOI: 10.1016/S0378-3758(96)00043-2. URL: https://www.sciencedirect.com/science/article/pii/S0378375896000432 (visited on 08/20/2024).

[2] David E. Tyler. “Statistical Analysis for the Angular Central Gaussian Distribution on the Sphere”. In: *Biometrika* 74.3 (Sept. 1987). Publisher: JSTOR, p. 579. ISSN: 00063444. DOI: 10.2307/2336697. (Visited on 10/21/2022).

[3] G.D. Forney. “The viterbi algorithm”. In: *Proceedings of the IEEE* 61.3 (Mar. 1973). Conference Name: Proceedings of the IEEE, pp. 268–278. ISSN: 1558-2256. DOI: 10.1109/PROC.1973.9030. URL: https://ieeexplore.ieee.org/document/1450960 (visited on 11/13/2024).

[4] Jesper Love Hinrich et al. “Archetypal Analysis for Modeling Multisubject fMRI Data”. In: *IEEE Journal on Selected Topics in Signal Processing* 10.7 (Oct. 2016). Publisher: Institute of Electrical and Electronics Engineers Inc., pp. 1160–1171. ISSN: 19324553. DOI: 10.1109/JSTSP.2016.2595103. (Visited on 01/24/2022).

[5] Alexander Schaefer et al. “Local-Global Parcellation of the Human Cerebral Cortex from Intrinsic Functional Connectivity MRI”. In: *Cerebral Cortex* 28.9 (Sept. 2018). Publisher: Cereb Cortex, pp. 3095–3114. ISSN: 1047-3211. DOI: 10.1093/cercor/bhx179. URL: https://pubmed.ncbi.nlm.nih.gov/28981612/ (visited on 09/08/2021).

[6] Ye Tian et al. “Topographic organization of the human subcortex unveiled with functional connectivity gradients”. en. In: *Nature Neuroscience* 23.11 (Nov. 2020). Publisher: Nature Publishing Group, pp. 1421–1432. ISSN: 1546-1726. DOI: 10.1038/s41593-020-00711-6. URL: https://www.nature.com/articles/s41593-020-00711-6 (visited on 05/15/2024).
